# Heteroskedasticity of neuronal population responses

**DOI:** 10.64898/2026.09.26.754693

**Authors:** Michael Okun

## Abstract

Typical changes in a neuronal circuit’s operating regime — such as shifts in sensorimotor conditions or in brain state — reshuffle firing rates across its neurons. One well-established key organising principle is rate preservation: firing rates correlate strongly across conditions. Here, we demonstrate that the joint distribution of firing log-rates across pairs of conditions exhibits an additional pervasive and previously unrecognised property: strong heteroskedasticity around the rate-preservation axis, whereby slow-firing neurons (< 1 spike/s) display substantially more variable relative responses across conditions than fast-firing neurons (>10 spikes/s). Using large-scale recordings across multiple brain regions (visual cortex, hippocampus, thalamus, and motor cortex) in mice and primates, we demonstrate that this structure is a generic feature of neuronal population responses. A spike subsampling procedure and explainable variance analysis confirm that this heteroskedasticity reflects true biophysical variance rather than statistical estimation bias resulting from low spike counts. Rate and spiking network models reproduce this phenomenon, revealing its key mechanism: while the standard neuronal transfer function (the *F*—*I* curve) is convex, its log-rate counterpart (the log*F*—*I* curve) is concave. Consequently, for equivalent input current shifts slow-firing neurons exhibit greater relative sensitivity whereas fast-firing neurons exhibit greater absolute sensitivity. Finally, we show that heteroskedasticity strongly influences downstream readout. Under wiring constraints, neurons with intermediate firing rates provide optimal discrimination performance, whereas under metabolic constraints, large assemblies of slow-firing neurons are most advantageous. These findings identify heteroskedasticity as a fundamental organising principle of neuronal population responses, establishing a mechanistic link between non-linear cellular transfer functions, population-level responses and energy-efficient representations.

## Introduction

Skedasticity is a basic property of multidimensional distributions. In the simplest 2-dimensional case, it characterises the variance of one variable along (conditioned on) the other variable (James *et al*., 2013). If the variance is uniform, as in the case of Gaussians, the distribution is said to be homoskedastic (Fig. 1A). Otherwise, the distribution is heteroskedastic (Fig. 1B). Skedasticity is an altogether separate property from correlation between the two variables (Fig. 1C) – intuitively, skedasticity to correlation is what variance is to mean in the case of one-dimensional distributions.

**Figure 1.**
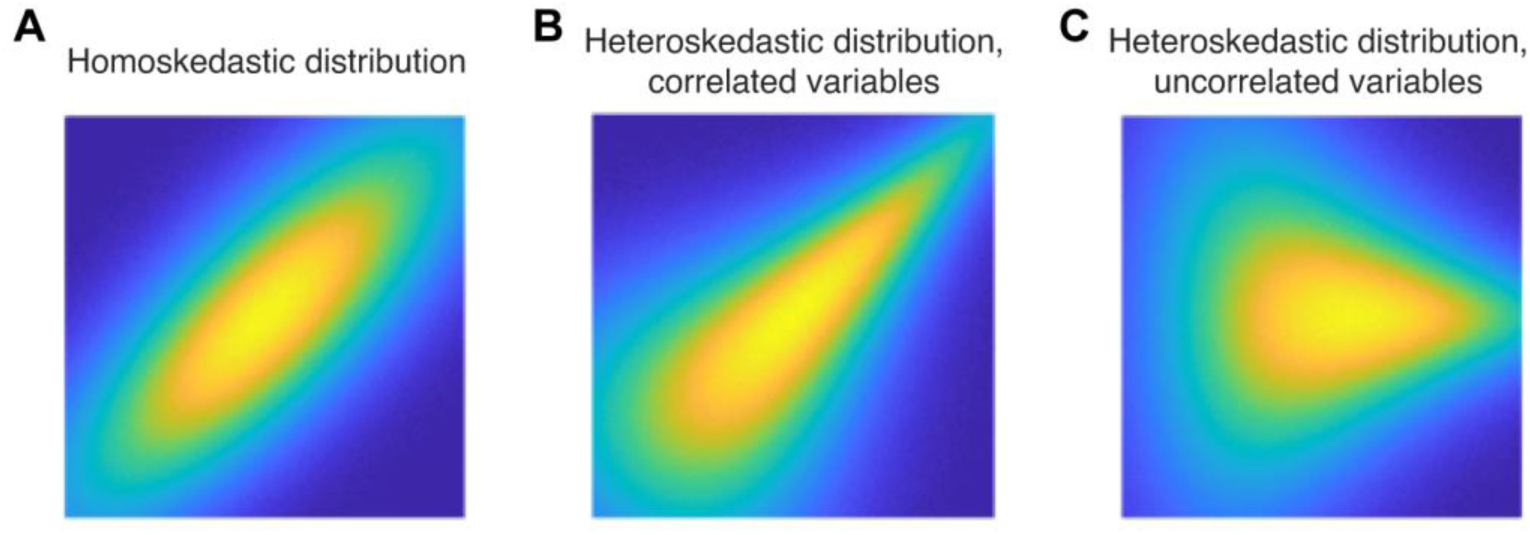
Homoskedasticity vs heteroskedasticity. **(A)** For a 2-dimensional Gaussian distribution (*X*, *Y*), the variance of *Y* conditioned on *X* = *x* is the same for all values of *x*, hence the distribution is homoskedastic. **(B)** Example heteroskedastic distribution (*X*, *Y*) in which *X* and *Y* are positively correlated. **(C)** Example heteroskedastic distribution in which *X* and *Y* are not correlated.

In (Dearnley *et al*., 2023; Jones, 2024) we demonstrated that the distribution of firing rates in pairs of brain states exhibits prominent heteroskedasticity, with much lower variability across high-rate neurons than across low-rate neurons, so that the joint distribution of neuronal log-rates in two brain states looks as Fig. 1B. This observation held for many kinds of brain state transitions and in multiple forebrain areas. Here, we extend this finding to demonstrate that this structure holds even more generally, applying to *any* typical change in a neuronal network’s operating regime. Such changes include, for example, transitions between the absence and presence of sensory or motor activity or between two distinct sensorimotor conditions. We use publicly available datasets from different brain regions of mice and monkeys to demonstrate this phenomenon. We then use generic recurrent rate and spiking network models to demonstrate that heteroskedasticity is explained by the nonlinearity of the neuronal transfer function. Finally, we demonstrate that heteroskedastic structure has a significant impact on the accuracy of downstream readout when the reader has wiring or metabolic constraints, with fast-firing neurons having poor performance in both cases (the term “fast-firing” should not be confused with “fast-spiking”, the former describes neurons with high firing rates, whereas the latter refers to neurons whose individual spikes are very short and produce a narrow waveform).

## Results

### Response heteroskedasticity

We start by considering the proverbial responses to gratings in the mouse visual cortex. We use a publicly available Neuropixels visual coding dataset from the Allen Institute (Siegle *et al*., 2021), in which awake head-fixed mice were presented with visual stimuli, while spiking activity in their left hemisphere was recorded with up to 6 Neuropixels probes. A “drifting grating” segment of these recordings contains many trials of responses to 4 stimuli (4 different drift directions, with the same contrast and spatial frequency; Fig. 2A) and is particularly suitable for our question.

**Figure 2.**
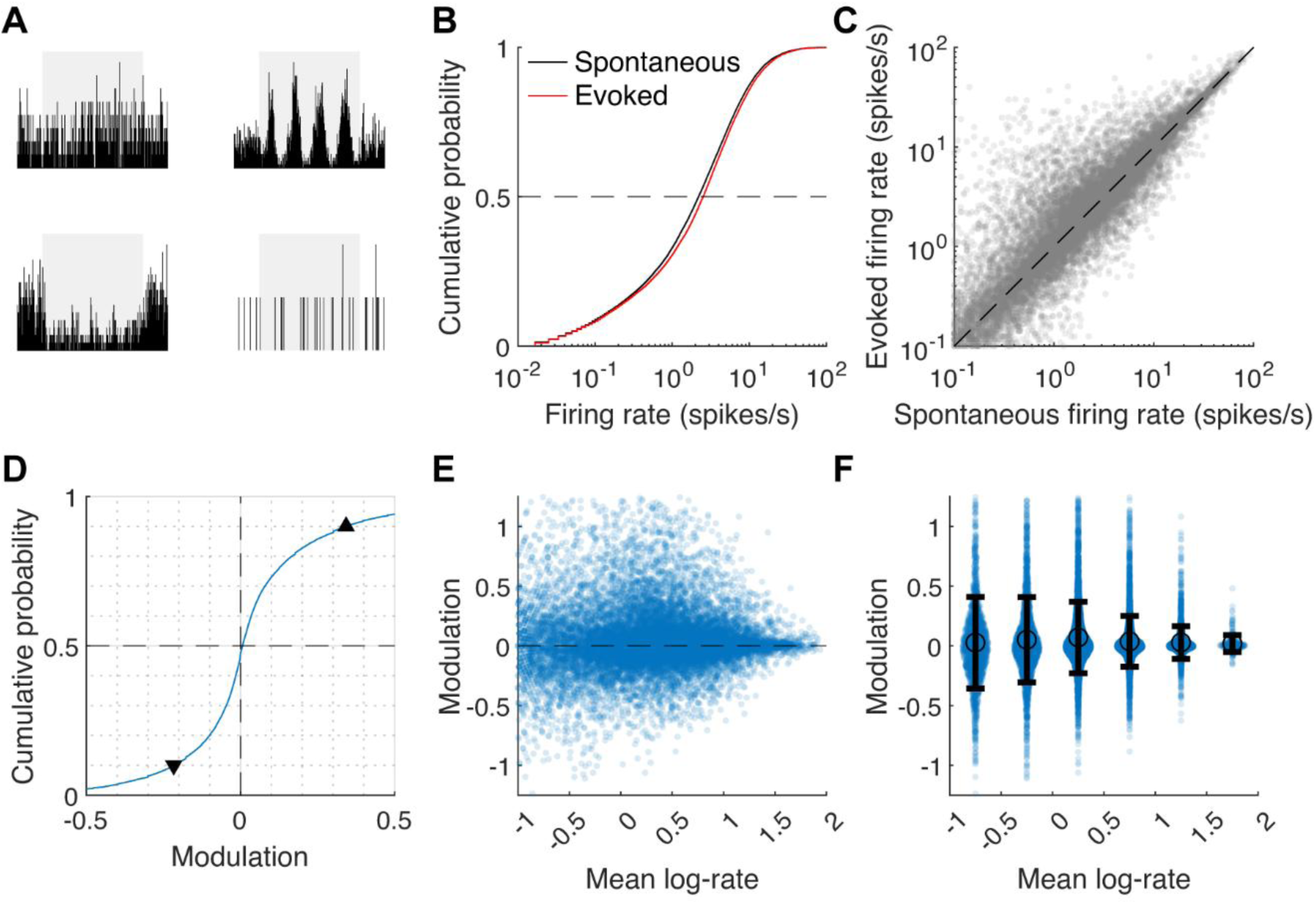
Heteroskedasticity of population responses. **(A)** Peristimulus time histograms (PSTHs) of four example neurons in visual cortex from one of the sessions in the Allen Institute dataset (responses to horizontal gratings), showing the evoked response (shaded area, 2s), with additional 0.5s on each side. **(B)** Cumulative distribution of the firing rates in spontaneous and driven conditions (n = 20285 neurons, 26 sessions). **(C)** Spontaneous firing rate vs rate during presentation of horizontal drifting grating of neurons in the visual cortex. **(D)** Cumulative distribution of rate modulation across these two conditions (▾ and ▴ indicate the 10^th^ and 90^th^ percentiles). **(E)** Modulation vs mean log-rate of each neuron, mathematically equivalent to (C) tilted 45 degrees right. **(F)** Same as (E) with neurons binned with half-decade spacing, with the mean and st.dev. of each bin shown – exhibiting clear heteroskedasticity of the distribution (cf. Fig. 1C).

The evoked rate of each neuron for a given direction was estimated from 60 trials. The spontaneous rate was estimated from inter-trial intervals (avoiding the offset responses) with the same total duration (see Methods for further details). The fact that evoked and spontaneous intervals are interleaved is important, since it allows to ascribe the differences in firing rates between the two conditions to the stimulus, rather than to slow activity changes, e.g., as a result of brain state changes.

Comparison of spontaneous and evoked firing rates across a large population of visual cortex neurons (20285 cells, 26 mice) leads to several observations. First, the change in the distribution of firing rates caused by visual stimulation is rather modest, e.g., the median rate increases from 2.4 to merely 2.7 spikes/s (Fig. 2B). Second, individual neurons have a strong tendency to preserve their rate (r = 0.92, p < 10^-100^, Spearman correlation; Fig. 2C), in agreement with previous suggestions (Buzsaki & Mizuseki, 2014). Third, neurons are split almost evenly between those that increase and decrease their rate. Quantitatively, the modulation, defined here as log_10_(*r_E_*/*r_Spont_*), where *r_E_* and *r_Spont_* are the evoked and spontaneous rates of each neuron, has population median that is very close to zero (Fig. 2D). It is however the case that modulation is stronger for upregulated neurons, e.g., its 10^th^ percentile is -0.22 but its 90^th^ percentile is +0.34 (▾ and ▴ in Fig. 2D), consistent with the increase in overall level of firing (Fig. 2B,C).

Visual inspection suggests that modulation is heteroskedastic, with slow-firing neurons further dispersed from the diagonal (Fig. 2C, cf. Fig. 1B; note that a neuron’s modulation, as defined here, is equal to its distance from the diagonal in Fig. 2C, up to a constant factor). This becomes apparent if we plot the modulation vs the position of each neuron on the firing log-rate spectrum (Fig. 2E; mean log-rate of a neuron is its average log-rate across the two conditions, namely log_10_(*r_E_* · *r_Spont_*)/2 = [log_10_(*r_E_*) + log_10_(*r_Spont_*)]/2, mathematically Fig. 2E is equivalent to rotation of Fig. 2C), and even more so if we bin the neurons by their mean log-rate (p < 10^-100^, Brown-Forsythe test for variance equality, Fig. 2F). It is also of some interest to note that modulation distribution is heavy-tailed (overall kurtosis of 12.5, and kurtosis of 6.5-12.9 in individual bins in Fig. 2F).

We will now proceed to demonstrate that the observed heteroskedasticity is a real neurophysiological effect, rather than a statistical estimation bias. Then we will show that it is not specific to mouse visual cortex but is a general phenomenon that can be observed in other brain areas as well as in primate recordings.

### Neuronal heteroskedasticity is not a statistical bias

The firing rate values for each condition in Fig. 2 come from trials whose combined duration is 2 minutes; hence these values are merely estimates of the “true firing rate” of the neurons in each condition. Within a 2-minute interval fast-firing neurons emit many hundreds of spikes, thus, their rate in each condition can be estimated rather precisely. At the same time slow-firing neurons emit a few dozen spikes or less, hence their rate estimate is unreliable. In other words, modulation estimate is noisier for slow neurons, and one might suspect that this explains the observation of heteroskedasticity in Fig. 2. For example, a simulation of a population of Poisson neurons with widely distributed baseline rates and true modulation that is independent of the baseline rate, shows that modulation estimated from 2 minutes of data is more widely distributed among the slow neurons (Fig. S1A). Even if there is no modulation at all, and we sample twice from Poisson neurons with the very same rates, we will find that modulation estimates exhibit heteroskedasticity (Fig. S1B).

To overcome this inherent bias in modulation variance estimation, we introduce the following approach. For a fixed threshold *N_T_*, we discard neurons which emitted less than *N_T_* spikes over the 4 minutes of activity (2 minutes × 2 conditions), whereas for any other neuron, *N_T_* spikes are picked at random from all those that were emitted. We use only these *N_T_* spikes to calculate a new modulation estimate log_10_(*N_E_*/*N_Spont_*), where *N_E_* and *N_Spont_* denote the total number of spikes in each condition after subsampling (*N_E_* + *N_Spont_* = *N_T_*). By construction, this estimate (to which we will henceforth refer as “fixed-sample modulation”) uses equal number of spikes for each neuron. In this manner the dependence of modulation estimate accuracy on firing rate, which exists if all the recorded spikes are used, is removed (see Methods for further details). In particular, when we apply this approach to the simulated Poisson data, no heteroskedasticity is observed (Fig. S1C,D).

Applying the fixed-sample approach to the drifting grating data reveals prominent heteroskedasticity. The st. dev. of fixed-sample modulation is 0.32 across neurons with mean log-rate below 0.5, and it drops to 0.13 across neurons whose mean log-rate is above 1.5 (p < 10^-80^, Fig. 3A). When the analysis was repeated for each recording session separately (the number of included neurons was 287-755), significant heteroskedasticity was observed in 23 of the 26 recordings. In the 3 remaining sessions, the modulation variance was also lower for fast-firing neurons, but this difference did not reach significance. Of note, fixed-sample modulation distribution was heavy-tailed, similarly to modulation (overall kurtosis of 10.8, and kurtosis of 4.1-10.9 in individual bins in Fig. 3A).

**Figure 3.**
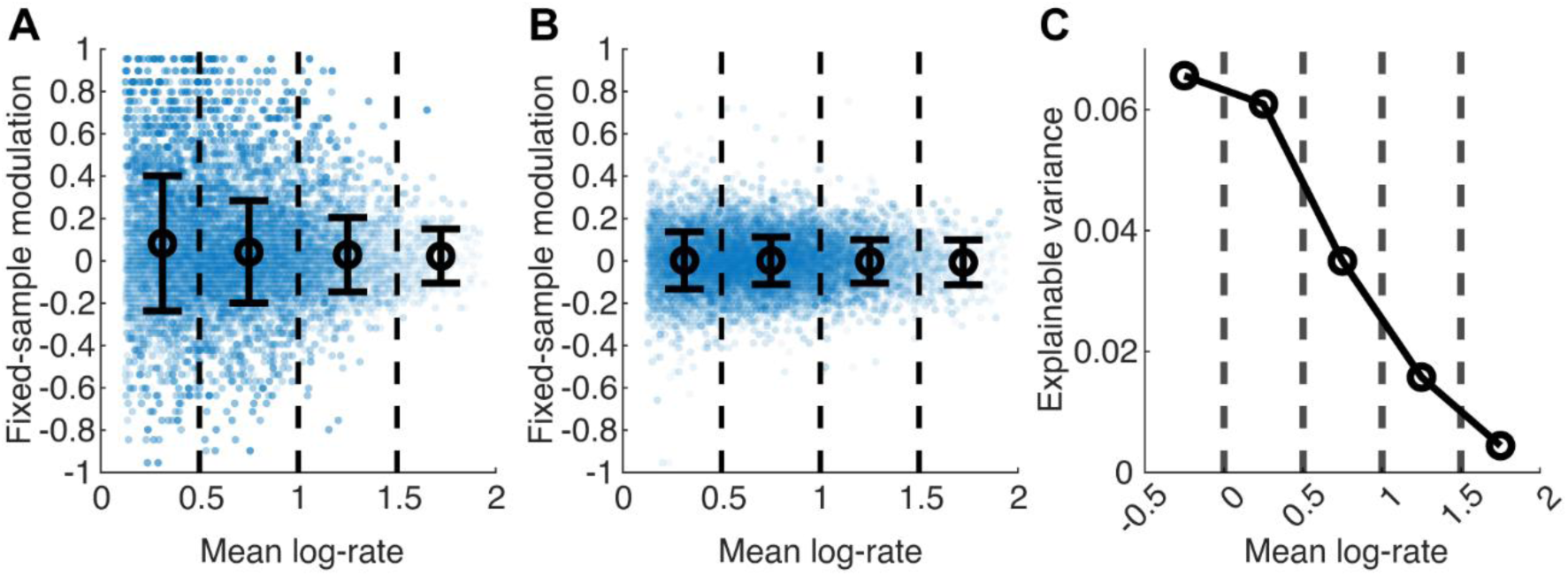
Physiological heteroskedasticity revealed by the subsampling method. **(A)** Fixed-sample modulation vs mean log-rate of each neuron (n = 12813 neurons), for spontaneous and drifting-grating conditions, showing clear heteroskedasticity (p < 10^-80^). **(B)** As in (A) when modulation is estimated across two random sets of trials in the evoked condition (p < 10^-35^, Brown-Forsythe test). (**C**) Explainable variance of modulation within each half-decade bin.

As a control, we also evaluated the modulation between two non-overlapping random subsets of trials within the same condition. In this case st. dev. had similar values in the different bins (between 0.13 and 0.1, Fig. 3B; we use 30 trials, equivalent to 60 s of activity, for both Fig. 3A and Fig. 3B). The small differences in the level of variance remained statistically significant, which likely reflects a true difference in within-condition variability between slow- and fast-firing subpopulations.

Next, we examined firing rate changes between two directions of drifting grating stimuli, rather than spontaneous vs stimulus conditions. Again, strong heteroskedasticity was present (fixed-sample modulation st. dev. falls from 0.30 to 0.12, p < 10^-75^, Fig. S2A; heteroskedasticity was significant in 22/26 sessions considered individually).

Finally, we verified the presence of heteroskedasticity using an altogether distinct statistical method. It involved the calculation of the explainable modulation variance, by using modulation estimated (without subsampling) from half of the trials as a predictor for modulation estimated from the other half (see Methods). In agreement with fixed-sample analysis, this approach also demonstrated that modulation variance is considerably lower among fast-firing neurons (Figs. 3C, S2B).

To sum up, there is clear evidence that heteroskedasticity is a true biophysical phenomenon in the mouse visual cortex: firing modulation produced by visual stimuli becomes progressively less variable across subpopulations of neurons with higher rates.

### Heteroskedasticity is a generic property of neuronal population responses

To examine how generic heteroskedasticity is, we tested for its presence beyond mouse neocortical areas. We started by examining modulation caused by drifting gratings vs spontaneous activity in thalamic lateral posterior nucleus (LP) and in the CA1 area of the hippocampus, as the Allen dataset includes a large number of neurons from these brain areas. In both cases, neuronal populations showed prominent heteroskedasticity (in LP st. dev. falls from 0.37 to 0.11, p < 10^-10^, 1553 neurons, Fig. 4A; in CA1 st. dev. falls from 0.30 to 0.12, p < 10^-50^, 10155 neurons, Fig. 4B). The case of the LP nucleus is of particular interest since, unlike cortical and CA1 circuits, it is a purely feedforward network lacking inhibitory neurons.

**Figure 4.**
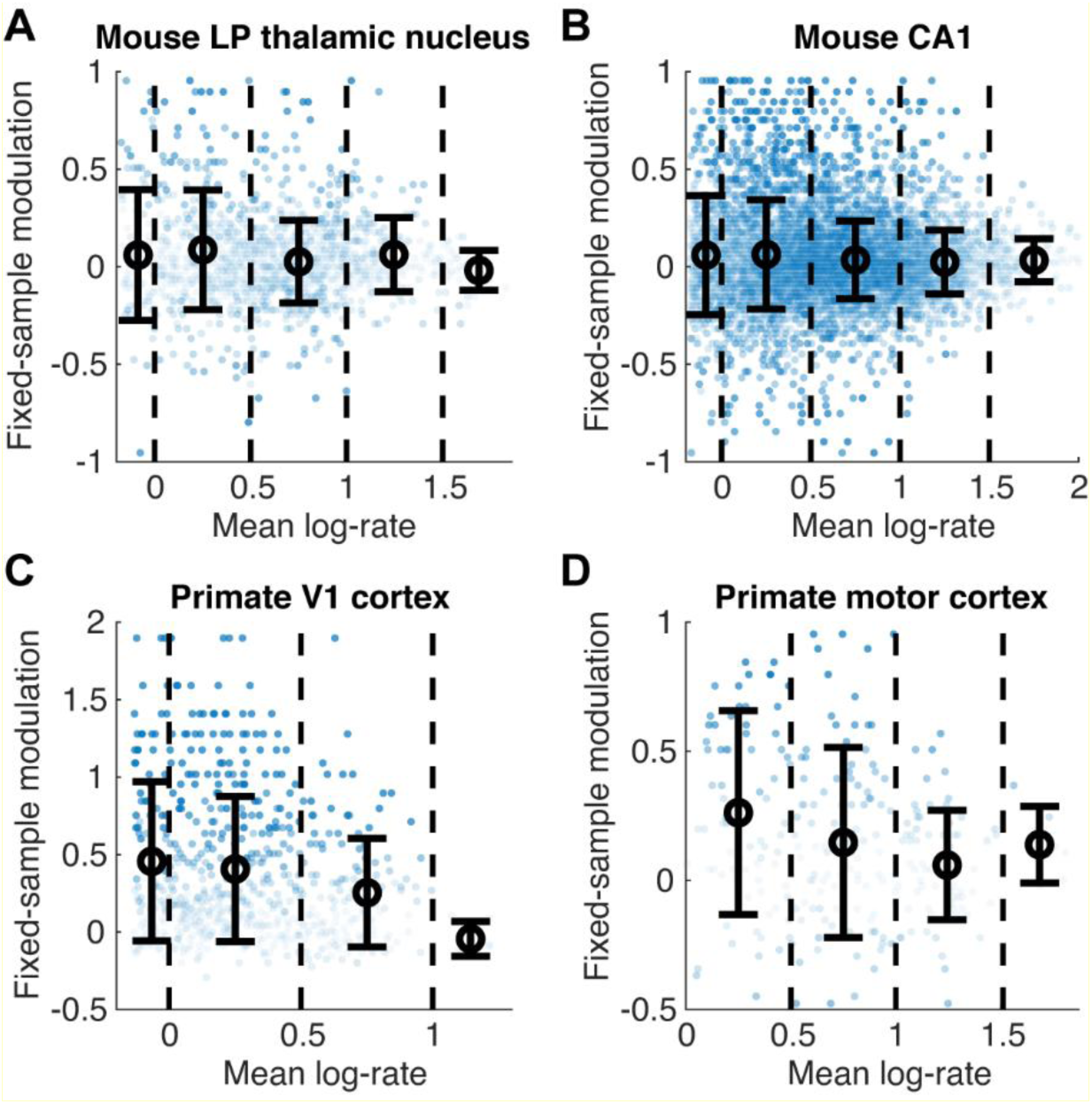
Heteroskedasticity across brain areas and species. **(A, B)** Heteroskedasticity of rate changes between spontaneous activity and gratings for neurons in thalamic nucleus LP and CA1 area of the hippocampus of the mouse (n = 1553, p < 10^-10^ and n = 10155, p < 10^-50^). **(C)** Heteroskedasticity of rate changes between spontaneous activity and visual stimulation (drifting gratings, multiple directions) for neurons in primate primary visual cortex (n = 650, p < 10^-4^). **(D)** Heteroskedasticity of rate changes upon movement initiation in primate motor cortex (n = 271, p < 5×10^-4^).

Next, we verified that heteroskedasticity is not unique to the mouse brain. First, we compared rate modulation between spontaneous activity and activity during drifting gratings presentation in primary visual cortex of anaesthetised monkeys (see Methods). We found prominent heteroskedasticity in this dataset (st. dev. falls from 0.56 to 0.11, p < 10^-4^, 650 neurons, Fig. 4C). We then considered the activity of neurons in primary motor and dorsal premotor areas (M1 and PMd) in two monkeys performing cued reaching movements (Lawlor *et al*., 2018). We compared rates around cue onset and the maximal velocity points. Again, we observed clear heteroskedasticity of rate modulation (st.dev. falls from 0.38 to 0.06, p < 5×10^-4^, 271 neurons, Fig. 4D).

We conclude that modulation heteroskedasticity is a general phenomenon occurring in a broad range of brain areas.

### Heteroskedasticity mechanisms

Observations of heteroskedasticity presented above, combined with our previous findings of heteroskedasticity across brain state transitions on timescales of minutes (Dearnley *et al*., 2023; Jones, 2024), suggest that it is a fundamental feature of neuronal responses. It is therefore plausible that response heteroskedasticity is present in basic models of neuronal circuits.

To test this hypothesis, we considered a rate network model of the cortical circuit. We opted for a simple “vanilla” version, with 4000 identical excitatory and 1000 identical inhibitory neurons, connected randomly with 10% probability and driven by constant external input (see Methods for further details). The network’s dynamics exhibits a chaotic, strongly coupled regime characteristic of cortical activity, with random fluctuations in neurons’ activity and a wide distribution of firing rates (Fig. 5A). To simulate a stimulus-evoked network state, we used an additional input to a random third of all the neurons. This input comprised a constant positive current whose magnitude was random for each cell (see Methods). Introduction of this additional input, or a switch between two such inputs, produced rate reshuffling, as reported previously and expected from theory (Sanzeni *et al*., 2023), and similarly to what we observed in the physiological data.

**Figure 5.**
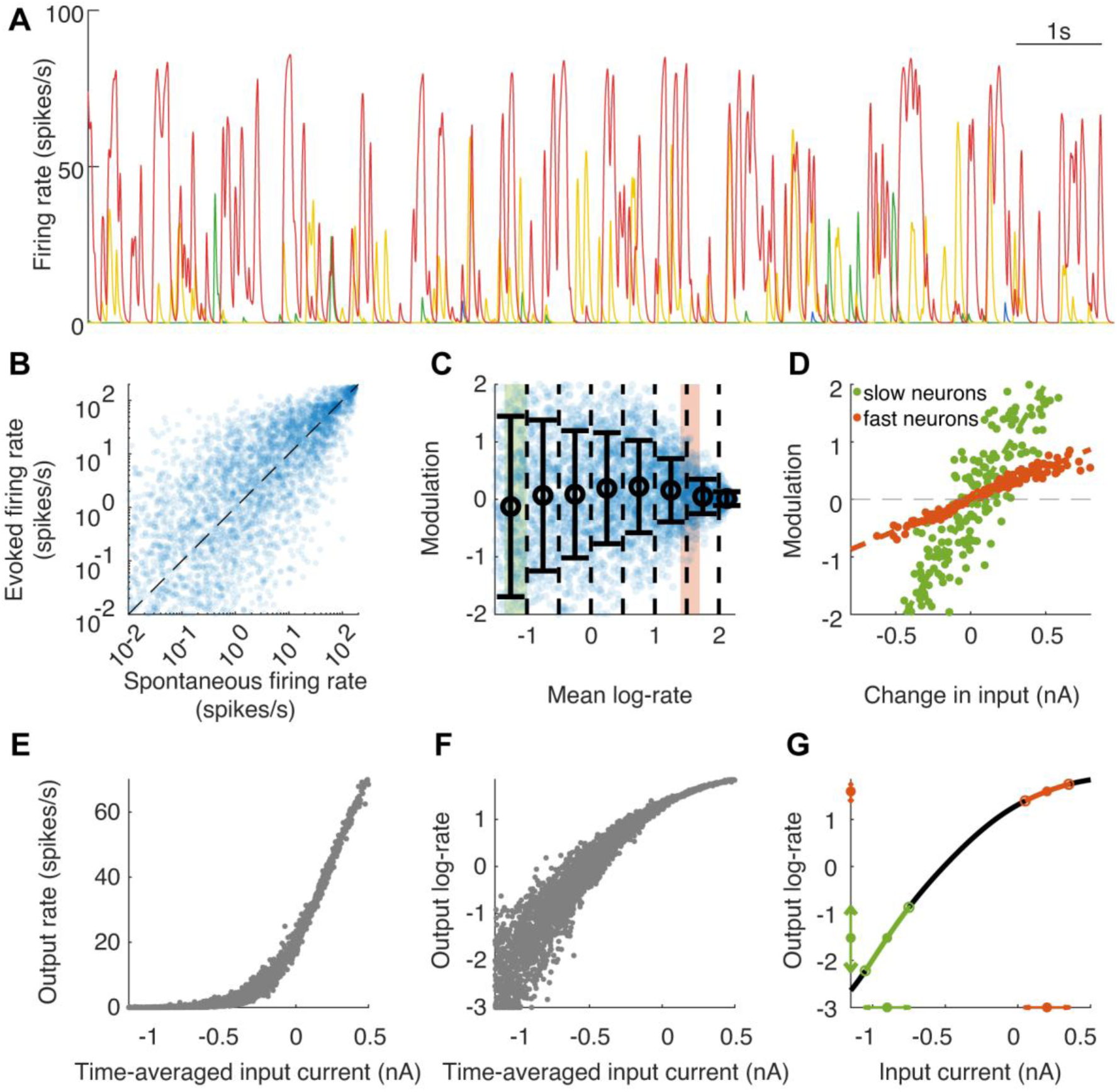
Heteroskedasticity in a rate network model. **(A)** Rates of four randomly selected neurons, illustrating the dynamics of rate fluctuations of individual neurons and the wide distribution of rates across neurons in the rate network model (blue, green, yellow and red, ordered by their average rate). **(B, C)** Heteroskedasticity of rate modulation between spontaneous and evoked regimes in the model. Shaded areas - log-rates of two example subpopulations of 200 E-neurons (slow and fast). **(D)** Change in the time-averaged input current caused by shift between spontaneous and evoked regimes has different modulation gain in slow- and fast-firing E-neurons. **(E)** The relationship between time-averaged input current and output rate for E-neurons in the model, notice the convex shape of the curve. **(F)** Same as (E), with output rates shown on log-scale, note the concave shape of the curve. **(G)** The average transfer function of E-neurons in the network. A change of ±0.15 nA in input (arrows along the x-axis) produces strong changes in log-rate of neurons on the left side of the curve (slow-firing neurons, green), but weak changes in log-rate of neurons on the right side of the curve (fast-firing neurons, brown).

Comparison of firing rates between the baseline and evoked conditions (Fig. 5B) revealed clear heteroskedasticity (st.dev. of modulation drops from 1.19 to 0.11, p < 10^-50^, Fig. 5C; cf. heteroskedasticity within the same condition, Fig. S3A,B). Heteroskedasticity was also prominent for rate changes between two distinct input patterns (Fig. S3C).

What is the source of heteroskedasticity in the model? To answer this question, let us focus exclusively on excitatory neurons (E-neurons), which comprise 80% of all the neurons. All E-neurons in the model are identical, hence any differences in their firing rates are fully determined by differences in input. We therefore examined how the change in time-averaged input between the baseline and evoked conditions translated into a change in output rate in subpopulations of slow-firing and fast-firing E-neurons. One example subpopulation was taken to include 200 E-neurons with mean log-rates between -1.35 and -1.0 (corresponding to 0.05-0.1 spikes/s) while the other included 200 E-neurons with log-rates between +1.4 and +1.7 (corresponding to 25-50 spikes/s). Within these two example subpopulations, the change in input current between the spontaneous and evoked conditions spanned approximately the same range, and was strongly correlated with the change in output log-rate (r = 0.94 and r = 0.98, p < 10^-80^ in both cases, Fig. 5D). However, the slope of the correlation between the change in input and modulation (the gain) was profoundly distinct in these two subpopulations (4.3, 95% CI [4.0, 4.5] vs 1.1, CI [1.05, 1.12]; Fig. 5D). In other words, the very same change in the time-averaged synaptic input results in a small difference in the log-rate of a fast-firing E-neuron but a large difference in the log-rate of a slow-firing E-neuron.

The reason for the vastly different gain between the fast and slow subpopulations becomes clear if we consider the effective F-I (frequency-current) transfer relationship that exists in the network. In typical *in vivo* conditions (the so-called fluctuation-driven regime) which are replicated by our rate network model, the F-I curve is convex (Fig. 5E and Fig. S3D; in full agreement with Roxin *et al*., 2011; Petersen & Berg, 2016). However, if the firing frequency is measured on a logarithmic rather than linear scale, to indicate the relative change in rate (henceforth we will call it “logF-I curve”), its relationship with input current becomes concave (Fig. 5F). Therefore, a change of equal magnitude in input current produces a much stronger change of log-rate in slow-firing neurons (on the left side of the logF-I curve) than in fast-firing neurons (on the right side of the logF-I curve; Fig. 5G).

Although the change in mean input current is the main factor driving heteroskedasticity, the analysis in Fig. 5 suggests that it is not the sole mechanism, because neurons with similar time-averaged input can differ quite substantially in their log-rate if they are slow-firing but not fast-firing (Fig. 5F). Furthermore, changes in input (across conditions) explain 96% of the changes in log-rate for fast neurons but only 88% for slow neurons (Fig. 5D). To explicitly test for such additional factors, we examined how changes in higher moments of the input relate to changes in log-rate modulation.

For input st.dev. the range of changes was comparable between our two example subpopulations, but the gain was substantially higher for slow-neurons (9.1, CI [7.6, 10.7] vs 0.3, CI [0.03, 0.6]; Fig. S3E). Furthermore, st. dev. explained significantly more of the residual modulation in slow-neurons than in fast-neurons (r = 0.64, p < 10^-20^ vs r = 0.15, p = 0.033). This difference is also due to the concave shape of the logF-I curve, since changes in fluctuations around the mean input have a stronger impact on the log-rate in the slow-firing neurons. Similarly, changes in input skewness explained residual modulation (after accounting for mean and st.dev.) in slow-firing neurons, but not fast-firing neurons (r = 0.46, p < 10^-10^ vs r = -0.13, p = 0.06; Fig. S3F). Input kurtosis changes were not correlated with residual modulation (after accounting for the first 3 moments) in either subpopulation (p = 0.19 and p = 0.6).

We expected the findings to generalise to conductance-based model of spiking neurons. To test this prediction, we implemented a simple “vanilla” version of a recurrent cortical network with 4000 identical excitatory and 1000 identical inhibitory leaky integrate and fire (LIF) neurons in a balanced state with irregular spiking activity, driven by constant and equal external input to all the neurons. As in the rate model, an additional input to a random third of neurons (a constant excitatory conductance) produced a perturbation of rates. These changes in firings rates exhibited clear heteroskedasticity and were explained by the same mechanisms identified in the rate network (Fig. S4).

The concave shape of the logF-I curve explains an additional observation, which we made in (Dearnley *et al*., 2021, 2023), having to do with the way log-rate distributions change between brain states in the experimental data. We consistently found that upon a change in state, the log-rate distribution of the upregulated neurons was squeezed (in addition to getting shifted rightward), whereas the log-rate distribution of the downregulated neurons was stretched (in addition to getting shifted leftward). The mechanistic explanation for the change in the width of these log-rate distributions is that neurons which are pushed rightwards change their log-rate progressively less, whereas neurons which are pushed leftwards change their log-rate progressively more (Fig. 5G).

To sum up, the rate and spiking models exhibit heteroskedasticity similarly to experimental data. Analysis of both models revealed that the main cause is that the neuronal input-output relationship, when firing rates are considered on logarithmic scale which reflects relative changes in firing (the logF-I curve), is concave. The fact that heteroskedasticity is dictated by the concave shape of logF-I relationship, which is shared by neurons across the nervous system, is consistent with the ubiquity of heteroskedasticity in neuronal population responses. The models also demonstrated that slow-neurons are more sensitive to changes in higher moments of the input than fast-firing cells, which also contributes to heteroskedasticity.

### Downstream decoding

We have established that response heteroskedasticity, with slow-firing cells exhibiting a proportionally stronger modulation, is a ubiquitous feature of neuronal responses. In some cases, e.g., for rate changes across prolonged brain states, such structure might not have functional significance in and of itself. However, when the conditions are informative for downstream brain areas, e.g. two drifting grating orientations in a visual two-alternative choice task, it is plausible that heteroskedasticity is important for the organisation of downstream readout.

To understand the consequences of the heteroskedastic form of the joint distribution of modulation and rate for readout, we examined the discrimination accuracy using the visual cortex drifting grating data from Figs. 2-3. Concretely, we examined how the probability of correctly distinguishing between two directions (here 45° vs 90°, the findings were similar for other direction pairs) changes along the firing rate spectrum for single neurons and for assemblies comprised of neurons with similar rates.

A priori, three key factors determine the accuracy of readout based on spike counts of individual neurons. First, the mean log-rate: a higher rate means a higher signal-to-noise (spike count mean to st. dev.) ratio, and thus higher accuracy. Second, the modulation strength: stronger modulation means higher accuracy. Third, the level of trial-to-trial variability, e.g., as quantified by the Fano factor (spike count variance to mean ratio), which is detrimental to accuracy. It is well established that most neurons exhibit supra-Poisson variability, which furthermore increases with rate (Goris *et al*., 2014).

We started by empirically estimating the spike count discrimination accuracy of each visual cortex neuron (by finding the optimal threshold for differentiating between a neuron’s responses to the two stimuli, see Methods for details). For neurons with very low rates, accuracy was just slightly above the chance level of 0.5 (Fig. 6A), and then rapidly increased with rate, so that among neurons with log-rates of ∼0 (i.e. rates of ∼1 spike/s) the best 10% had accuracy values > 0.7. However, beyond this point accuracy plateaued: its values for neurons with log-rates of ∼1.5 (corresponding to rates of > 30 spikes/s) were not discernibly higher. These observations suggest that as one moves right along the firing rate spectrum, at first the gain from having more spikes dominates and pushes the accuracy up, but from the midway point further accuracy gain is prevented by increasingly weak modulation and higher Fano factors.

**Figure 6.**
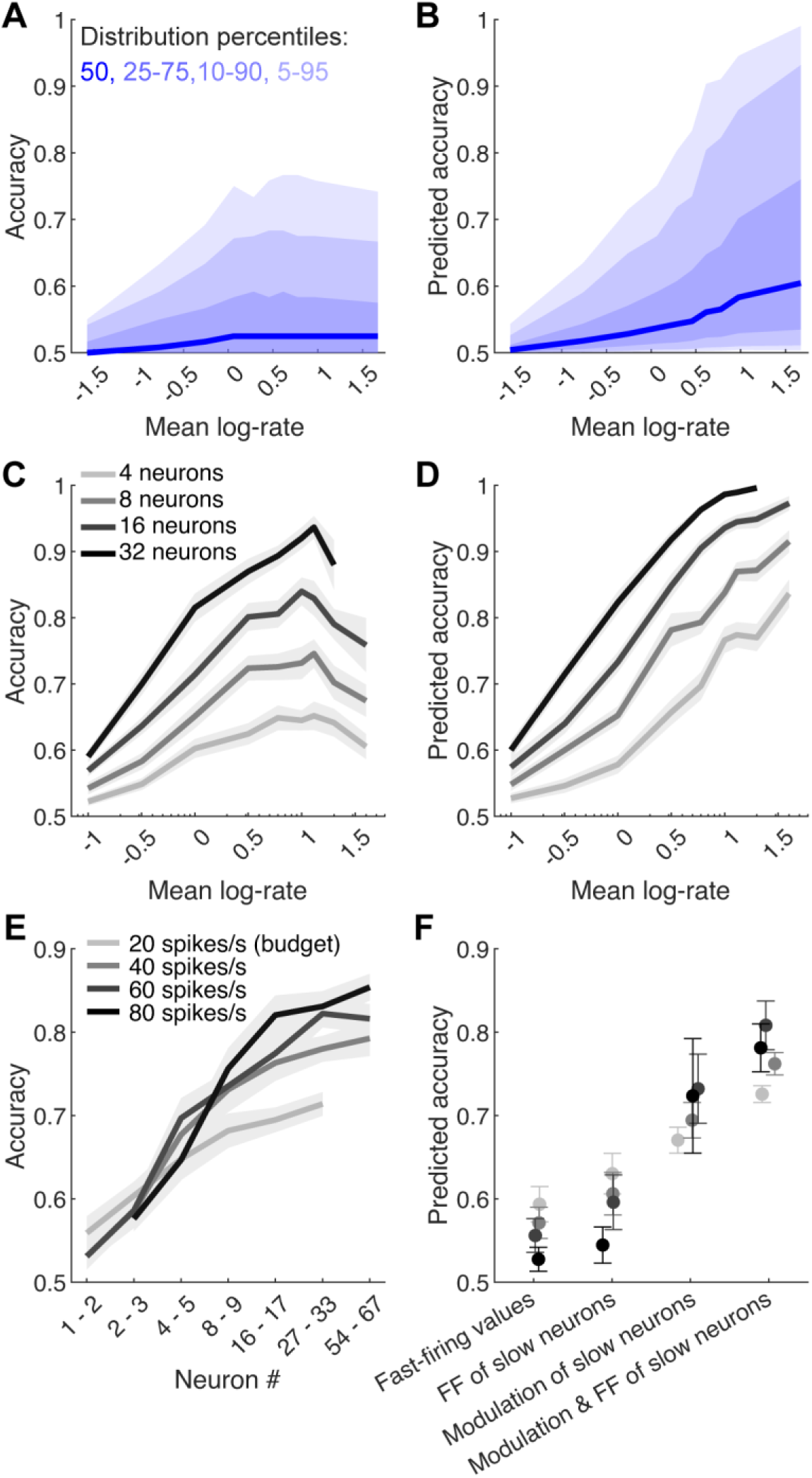
Heteroskedasticity’s impact on readout accuracy. **(A)** Distribution of discrimination accuracy vs mean log-rate of individual neurons in the mouse visual cortex – empirical data. Different shades show the median, 25-75, 10-90 and 5-95 percentiles. **(B)** As in (A), using analytically calculated accuracy in a synthetic population whose statistics match the empirical data, except for modulation heteroskedasticity, which does not exist (modulation strength is independent of neurons’ location on the rate spectrum). **(C)** Discrimination accuracy using spike counts from randomly selected assemblies of neurons (of size 4-32) with similar mean log-rates spanning the rate spectrum. **(D)** As in (C), in a synthetic population whose statistics match the empirical data, except for modulation heteroskedasticity, which does not exist. **(E)** Accuracy of randomly selected assemblies of different sizes conforming to a fixed total spike budget – empirical data. **(F)** Analytically calculated accuracy of a single neuron with spike budget of 20-80 spikes/s in four cases: (i) its modulation and Fano factor (FF) parameters are median modulation and FF values of neurons whose rate matches the budget (hence accuracy closely matches that of assemblies comprised of just one neuron in E), (ii) modulation parameter as before, FF parameter is the median of FF of neurons with log-rates of ∼0, (iii) FF parameter as in (i), modulation parameter is the median modulation strength of neurons with log-rates of ∼0, (iv) both modulation and FF parameters are medians of the slow neurons. In (C-F) shaded area and error bars show SEM, based on averages over n = 26 data points corresponding to the recording sessions.

To test this explanation, we analytically predicted the accuracy of each neuron from the three key parameters determining accuracy (see Methods for details). This prediction closely matched the empirical accuracy values (Fig. S5A,B). The analytical model therefore allowed us to determine how accuracy would have behaved if modulation strength of each neuron was independent from its position on the rate spectrum; we did so by reshuffling the modulation strength parameter across the neuronal population before applying the prediction. In this scenario, accuracy increased across the entire rate spectrum, rather than just its first half (Fig. 6B, cf. Fig. 6A). We used the model in a similar manner to quantify the effect of Fano factor’s dependence on rate. This effect was much weaker than the effect of modulation strength decrease (Fig. S5C,D). We conclude that Heteroskedasticity structure has a dominant effect on accuracy of individual neurons.

Next, we turned to examine the accuracy of readout from assemblies of neurons. If there are no constraints on the assembly, then all of the visual cortex neurons can be utilised, leading to superb performance (Stringer *et al*., 2021). However, such readout is not biophysically realistic, since actual readout in the brain is highly constrained – “information costs energy and space” (Sterling & Laughlin, 2015).

To remain within the biophysically relevant realm, we started by considering the case of a downstream reader whose number of inputs is capped. Specifically, we examined the discrimination accuracy of sets of 4, 8, 16 and 32 neurons. We populated each set by simultaneously recorded neurons with similar mean log-rates (allowing differences of at most 0.25, see Methods). Discrimination was performed based on the spike count vectors using a simple linear readout (linear support vector machine, e.g. as in Graf *et al*., 2011, see Methods for further technicalities). Similarly to the case of single neurons, and for all assembly sizes, accuracy rose with rate at first, but then flatlined and even declined (Fig. 6C).

To understand this behaviour, we first noted that noise correlations did not play an important role, since their removal (by shuffling trials across neurons) had minimal effect on accuracy (the values after removal of noise correlations explained 94% of the original accuracy values, Fig. S6A). This in turn allowed us to use a simple computational model to predict an assembly’s accuracy using the mean log-rate, modulation strength and Fano-factor of each neuron (Fig. S6B,C; see Methods for details). As in the case of single neuron accuracy, reshuffling the modulation values across the neuronal population revealed that accuracy would have kept increasing if it were not for the weak modulation of the high-rate neurons (Fig. 6D). The contribution of Fano-factor’s dependence on rate was small (Fig. S6D,E). This demonstrates that, just as in the case of single neurons, heteroskedasticity structure has a dominant effect on readout accuracy of neuronal assemblies with constrained bandwidth.

Next, we considered the case of a downstream reader whose metabolic cost, rather than input bandwidth, is limited. To examine the key implications of such kind of constraints without going into detailed modelling of metabolic processes, we will presume that the downstream reader has a capped spike budget, e.g., a total input rate of 20 spike/s (on average across the two stimuli). In the case of independent Poisson neurons, the discrimination accuracy of one neuron which uses up all the spike budget is equivalent to that of a homogeneous assembly of neurons with the same modulation strength (e.g., a neuron with rates of 30 and 10 spikes/s in the two conditions provides the same accuracy as an assembly of 10 neurons each of which responds with rates of 3 and 1 spikes/s); this is an easily verifiable mathematical statement. To the extent that an independent Poisson population approximates how assemblies of real neurons behave, we expect the accuracy for a given budget to be similar for assemblies of different sizes. On the other hand, we have seen that the Poisson approximation is violated in two systematic ways. First, strong modulation is far less common among fast-firing than slow-firing cells, and second, the slower neurons have lower Fano-factors. Both of these factors suggest that for a given budget, an average large assembly of slow-firing cells provides a higher accuracy than an average small assembly of fast-firing cells.

To test this hypothesis, we performed the accuracy analysis for random assemblies of visual cortex neurons with similar log-rates (discrepancy of at most 0.15), with total output rate of the assembly capped at 20, 40, 60 or 80 spikes/s. We consistently observed that larger assemblies of slower neurons allowed for a more accurate readout, with assemblies of 1-3 cells having a typical accuracy of ∼0.6, whereas assemblies of ∼30-60 neurons achieved accuracies of 0.7 – 0.8 with the same spike budget (Fig. 6E).

In order to confirm that the higher accuracy of larger assemblies is explained by modulation difference along the rate spectrum, we cannot compare against a synthetic population in which modulation and log-rate are independent, as we have done above, because such synthetic population would have a higher metabolic cost, the very same quantity we are examining. (This is the case because a fixed mean log-rate does not translate into a fixed spike budget; stronger modulation requires more spikes). Instead, we analytically calculated the accuracy of a single neuron with spike budgets of 20-80 spikes/s, whose Fano factor, modulation strength or both were either assigned the median value empirically observed in neurons with rates of 20-80 spikes/s, or the median value observed in slow-firing neurons (with rates of 0.5 – 1 spikes/s, corresponding to rates of neurons in our large assemblies). This calculation confirmed that modulation strength and Fano factor explain the higher accuracy of larger assemblies, with modulation having the dominant role (Fig. 6F).

We have repeated all the analyses in Fig. 6 for CA1, since the Allen recordings included large samples of neurons in this area as well, and observed similar results, except that the accuracy values were somewhat lower than in the visual cortex (Fig. S7).

## Discussion

We analysed changes in firing rates driven by sensorimotor events in populations of neurons in cortical, hippocampal and thalamic areas of mice and monkeys. In all the examined cases, substantial heteroskedasticity of firing log-rate changes was observed (Figs. 1-4), with slow-firing subpopulation (rates below 1 spike/s) exhibiting modulation that was 2.5-6 times more variable than the fast-firing subpopulation (rates of 10-30 spikes/s and above). This behaviour was also found in off-the-shelf rate and spiking neuronal network models (Fig. 5), which revealed that the primary mechanism behind heteroskedasticity of log-rate responses is the concave shape of the transformation of input current into firing log-rate of neurons (logF-I curve). The fact that F-I curve is convex whereas logF-I curve is concave means that for a given fixed change in input, a fast-firing neuron is more sensitive in absolute terms (the change in the number of spikes produced every second), whereas a slow-firing neuron is more sensitive in relative terms (the proportional change). We showed that for the purpose of downstream readout in a limited bandwidth (number of inputs) scenario assemblies comprised of neurons with intermediate rates provide an optimal discrimination accuracy by balancing the number of spikes with reasonably strong modulation; for a reader with a limited spike budget (a metabolic energy constraint), large assemblies of slow-firing neurons are optimal (Fig. 6).

It is presently understood that neurons operate in two different regimes: the fluctuation-driven regime, in which the time-averaged input is below the spiking threshold of the neuron, and the mean-driven regime in which the input is strong enough to depolarise the membrane above the threshold. In the former case the spikes are primarily driven by occasional depolarising input transients which are strong enough to bring the membrane to the threshold, whereas in the latter case the neuron is constantly firing with a rate that depends both on the input and on the properties of the spiking mechanisms, such as after-hyperpolarisation currents and adaptation.

Spiking neurons adhering to Dale’s principle and receiving balanced excitatory and inhibitory inputs comprise a basic and evolutionary ancient network architecture throughout the central nervous system, irrespective of whether the network is recurrent or feedforward only (Levenstein & Okun, 2023). The fundamentals of operation of such networks are best understood through statistical physics perspective, showing that in these networks the fluctuation-driven regime dominates and the distribution of firing rates across neurons is necessarily broad (Roxin *et al*., 2011). Intuitively, just as one cannot have a gas with only slow or only fast molecules, one cannot have a network of only slow- or only fast-firing neurons.

The input-output transformation, also known as the F-I transfer function or curve, is a key concept in quantitative descriptions of neuronal activity, central to our understanding of the operating regime of neuronal networks. The input is typically specified by the synaptic current or by the resulting membrane voltage, and the output is given by the firing rate. The transformation was extensively investigated, both experimentally and computationally (McCormick *et al*., 1985; Brunel, 2000; Anderson *et al*., 2000; Miller & Troyer, 2002; Hansel & Vreeswijk, 2002; Chance *et al*., 2002; Priebe & Ferster, 2008; Silver, 2010; Roxin *et al*., 2011; Gerstner *et al*., 2014; Petersen & Berg, 2016; Levenstein *et al*., 2017; to name just a few of the key publications from a much broader literature). In the mean-driven regime the F-I relationship is approximately linear, followed by saturation for very high inputs which bring the spike generation to its limits. This saturating portion is not reached in our simulations (Figs. 5E, S3D, S4B) and is rarely reached *in vivo*. In the fluctuation-driven regime the F-I curve is convex, and closely follows a power law, i.e., *F* ∝ [*I*]*^p^* (*p* > 1). It therefore follows that in both mean-driven and fluctuation-driven regimes, log-rate is a concave function of the input (see also Fig. 3 in Levenstein & Okun, 2023).

Our modelling analysis (Figs. 5, S3, S4) demonstrated that in the presence of broadly distributed firing rates, response heteroskedasticity is an inevitable consequence. In this analysis we deliberately considered networks with identical excitatory and inhibitory cells, whereas neurons in brain circuits are highly heterogeneous. Indeed, dozens of genetically defined cell-types have been established even within individual brain areas (Tasic *et al*., 2018; Zhang *et al*., 2023). Cell-type diversity is likely to be an important part of the explanation of heteroskedasticity and of the broad distribution of firing rates observed *in vivo*. For example, in the mouse barrel cortex there is a ∼50-fold difference between median *in vivo* firing rate of layer 2/3 and layer 5 cells, which is clearly related to the different cell-types and connectivity of these layers (O’Connor *et al*., 2010), and interneurons have higher rates than excitatory cells (Gentet *et al*., 2012; Buzsaki & Mizuseki, 2014; Brecier *et al*., 2022; Valero *et al*., 2025). However, cell-type differences are unlikely to fully explain firing rate diversity and heteroskedasticity for the simple reason that these phenomena persist within more homogeneous neuronal subpopulations. For instance, firing rates are spread over many orders of magnitude even within layer 2/3 or layer 5 (O’Connor *et al*., 2010; Buzsaki & Mizuseki, 2014). Similarly, broad distribution of rates and heteroskedasticity across brain state transitions was observed in the population of narrow-spiking, putative inhibitory neurons (Dearnley *et al*., 2023). And if we consider the three main cortical interneuron subtypes (parvalbumin, somatostatin and vasoactive intestinal polypeptide) in isolation, for each subtype we again find neurons whose firing rates are spread over at least 1.5-2 orders of magnitude (Brecier *et al*., 2022). Therefore, we suggest that connectivity and input heterogeneity matter for firing rate statistics of physiological neural circuits as much as they do for models with homogenous intrinsic cell properties. Experimental studies that directly examined such difference between neurons with different rates support this suggestion (Yassin *et al*., 2010; Trojanowski *et al*., 2021).

The key insight that the F-I curve is convex whereas logF-I curve is concave implies that fast-firing neurons are more sensitive to input changes in absolute terms, whereas the slow-firing neurons are more sensitive in relative terms. In the final part of the present work, we argued that it is the relative sensitivity which matters for a realistic downstream readout. Specifically, we examined how well a downstream reader can discriminate between the two network conditions. A cell that is silent in one condition and has a very high rate in the other would be an ideal input source for such readout, however the heteroskedastic structure of modulation directly implies that such neurons are very rare or even non-existent. For a realistic neuronal population, the best input to a reader with a limited input bandwidth comes from neurons with intermediate rates of ∼1 spike/s and not the higher rate neurons, because the former have an optimal balance of number of spikes with sensitivity. For a reader with a limited metabolic (energy) budget, the bias towards slow-firing but larger assemblies was even more pronounced. This scenario provided a clear illustration of the advantage of readout from slow-firing neurons: individual slow neurons are significantly more sensitive (in relative terms), whereas assemblies comprised of many dozens of such cells generate enough spikes for a high signal-to-noise ratio and therefore high accuracy. This analysis therefore provides a mechanistically-informed explanation of the utility of the slow neurons (often 30% - 50% of the total neuron number) within brain circuits.

The readout analyses used a behaviourally relevant timescale of 2s, which was the duration of individual trials in Allen dataset. For decoding on a significantly shorter timescale (mice can discriminate 45° vs 90° drifting gratings within less than 100ms, Resulaj *et al*., 2018), fast-firing neurons are more advantageous than the slower neurons. This is consistent with the general idea that fast-firing neurons are more central to the networks’ operation and carry more basic information, whereas the slow-firing cells carry more refined signals which require longer processing times. It was suggested that the reason for this difference is saturation of the strength of synaptic inputs on the fast-firing neurons (Buzsaki, 2019). The mechanistic insights of the present work suggest a more nuanced interpretation, showing that the high rate and its stability across different regimes do not necessitate saturation, as was the case in our computational models.

The assembly sizes that were used in our analysis are compatible with our present understanding of communication between brain areas. For example, several different approaches indicate that layer 4 neurons in primary sensory cortices receive input from one-to-several hundred efferents from primary thalamic nuclei (Bruno & Sakmann, 2006; Chen *et al*., 2026). A recent brain-wide mapping effort shows that, just as one would expect, these thalamocortical connections are several-fold stronger than other inputs (Yao *et al*., 2023). That is, for a typical pathway, a neuron receives inputs from tens of presynaptic cells, just as in our analysis.

The assemblies we used are not fully realistic in that they were populated using all the visual cortex neurons recorded in a session, i.e., they could be recorded by different probes and thus be far away from each other. However, since the information on drift direction is present throughout the visual cortex, we believe that this limitation does not invalidate our findings. In the coming years, technological advances, such as fast, single-spike resolution imaging afforded by the most advanced voltage sensors, will allow collecting data which does not impose such constraints.

## Methods

### Datasets

#### Mouse data

Responses to drifting gratings are part of the publicly available Neuropixels visual coding dataset from the Allen Institute. Briefly, mice were implanted with headplates and habituated to being head-fixed on a running wheel. Recordings were performed with up to 6 Neuropixels 1.0 probes (Jun *et al*., 2017) inserted into primary and higher visual cortical areas, and also reaching down into the hippocampus and the thalamus. The recordings were processed via a robust pipeline that included spike-sorting with subsequent quality control. The positioning of the probes (that were covered by a fluorescent dye) was histologically verified, which allowed to establish the brain area for the vast majority of recorded neurons. Full details on this dataset are available in (Siegle *et al*., 2021), the accompanying technical white paper, and Allen Brain Atlas SDK (allensdk.readthedocs.io/en/latest/visual_coding_neuropixels.html).

Recording sessions lasted ∼3.5h and included several sets of stimuli. Here, we used one specific set, which included drifting gratings in 4 orientations (0°, 45°, 90° and 135°, with just one contrast and one temporal frequency of 2 Hz). These 4 stimuli (each trial was 2s in duration) were presented in a randomly generated order, for 300 trials in total and with 1s inter-trial interval. Here we used 0.5s interval preceding each trial as baseline (spontaneous) activity, thus avoiding offset responses in the 0.5s immediately following grating presentations.

#### Primate visual cortex data

We have used a dataset of recordings in macaque primary visual cortex. Briefly, prototype Neuropixels 1.0 probes were used to record the spiking activity across the layers of the cortex in 2 anaesthetised monkeys viewing drifting gratings. The dataset includes 5 sessions (3 sessions in monkey M1, 2 sessions in monkey M2), all of which were used for the analysis.

In each session we used 150 ms of activity preceding the stimulus onset as the first (spontaneous) condition, and 150 ms around the peak evoked population activity as the second (evoked) condition. Each session had 720 or 1440 or 2160 trials. For pooled analysis we used 720 trials (the minimum number of trials available across all sessions), for individual session analysis we used all the trials available for that session. Full details on this dataset are available in (Zhu *et al*., 2020; Trepka *et al*., 2022) and doi.org/10.5061/dryad.x3ffbg7p2.

#### Primate motor cortex data

We have used the pmd-1 dataset from the crcns.org repository (Lawlor *et al*., 2018). Briefly, neurons were recorded in primary motor cortex (M1) and dorsal premotor cortex (PMd) of two monkeys performing sequential reaching task. The animals controlled the position of a cursor on a computer screen using a manipulandum which could move in a horizontal plane. To obtain rewards monkeys had to perform reaching movements. The dataset includes 4 sessions (3 in monkey MT, and 1 in monkey MM).

In each session we used 415 reaches (the minimum number of eligible reaches available across all four sessions). The activity in a 200 ms interval around cue onset (-50 to +150 ms), during which the velocity of the cursor was minimal, was used as the first condition. The second condition was the spiking activity around peak cursor velocity (+300 to +500 ms from cue onset). Neurons from M1 and PMd were pooled for the purposes of the analysis. The number of neurons was too low to allow the analysis of individual sessions (as we did for other datasets), instead neurons were pooled across the sessions. Full details on this dataset are available in (Perich *et al*., 2018).

### Comparison of variance

Brown-Forsythe test (MATLAB’s vartestn function) was used to compare the variance of modulation values across half-decade bins to which individual neurons were assigned based on their mean log-rate (e.g. as seen in Fig. 3A). Throughout the text p-values presented without explicit mention of a statistical test are the results of such Brown-Forsythe test.

In scatter plots showing individual neurons (e.g. as in Fig. 3A), a few points might have fallen outside the axes limits and are not visible. The statistical tests included all data, irrespective of their visibility in the presented plots.

### Fixed-sample modulation

The main idea behind the fixed-sample procedure was outlined in Results. To recapitulate, neurons with < *N_T_* spikes across all the trials of the two conditions are removed. For any remaining neuron that emitted *N* ≥ *N_T_* spikes, *N* − *N_T_* spikes are picked at random (disregarding the condition to which any spike belongs) and removed, the modulation is calculated using the remaining *N_T_* spikes. In this manner modulation is estimated from equal number of spikes for all the neurons, and it is no longer the case that the estimate is noisier for slow-firing neurons. The value *N_T_* = 80 was used throughout.

The method has an additional caveat having to do with the fact that the procedure forces all neurons to have the same number of spikes after subsampling, whereas heteroskedasticity is quantified with firing rates on the logarithmic scale. This caveat is illustrated in Fig. S8, which reuses the data from Fig. 2, with spike counts from 4 minutes of recording (2 minutes of spontaneous activity and 2 minutes of drifting gratings; Fig. S8A reproduces Fig. 2E). After applying the subsampling procedure we get the data shown in Fig. S8B. Note that a neuron with a mean log-rate of -1 with spontaneous log-rate of -2 and evoked log-rate of 0 (i.e. a neuron that undergoes a very strong modulation resulting in a 100-fold difference in rate) is not removed (its rate of 1 spike/s in the presence of stimulus means that it has ∼120 spikes in total during 2 minutes of stimulation, well above the *N_T_* = 80 threshold). On the other hand, a neuron with mean log-rate of -1 that is not modulated by stimulus (i.e. it has a rate of 0.1 spikes/s in both conditions) would emit ∼12 spikes per condition (∼24 spikes in total over the 4 minutes, well below the *N_T_* = 80 threshold) and is excluded.

Since the bins are based on mean log-rates we must exclude neurons with *N_T_* spikes but low mean log-rate (neurons that fall in the area between the red dashed curve and vertical line in Fig. S8B,C). This means excluding neurons not only if they emit less than *N_T_* spikes but also if their mean log-rate is below log_10_(*N_T_*/T), where T is the total duration of the data used for the analysis (in the present example T = 240 s).

Finally, it should be noted that the subsampling procedure itself introduces non-uniform variability into the modulation estimate. This can be seen if we take the firing rates and the spike counts to be exactly equal across the two conditions for all the neurons. Although in this synthetic case the true modulation of each neuron is 0 (by definition), upon subsampling (which removes random subsets of spikes in each case), we will get non-zero modulation estimates, which furthermore will display heteroskedasticity. This heteroskedasticity however is small, and (more importantly) in reverse direction from the experimentally observed one - namely, the variability among the slow-firing neurons is lower as they have less spikes removed (st. dev. grows from 0.06 to 0.10, Fig. S8D).

### Explainable variance estimate

We partition the set of trials in each one of the two conditions (e.g. evoked and spontaneous activity) into two random disjoint subsets of equal size. Then, we calculate the modulation twice, each time using a different trial subset, i.e., for every neuron *i* we get two modulation estimates: 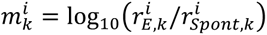, where 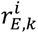 and 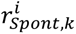 are the firing rates of the neuron in the two conditions in subset *k* (*k* = 1,2).

For a subset S of neurons, the explainable variance of modulation is [*SS_tot_* − *SS_err_*]_+_/|S|, where 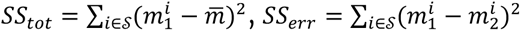, and *m̅* is the average modulation of the neurons in S in the first subset of trials. In our case the subsets S are all neurons with mean log-rate between 1.5 and 2, between 1.0 and 1.5, etc.

### Rate network model

We simulated a recurrent network of homogeneous excitatory (E) and inhibitory (I) neurons. We used a rate-based formalism, as described in multiple previous sources (Dayan & Abbott, 2005; Ahmadian *et al*., 2013; Sanzeni *et al*., 2023).

Briefly, the network has *N_E_* excitatory and *N_I_* inhibitory neurons, for a total of *N* = *N_E_* + *N_I_* neurons. At every time point *t*, the state of the network is defined by 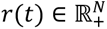, a vector of non-negative rates. The input received by the neurons is given by *u*(*t*) = *W* · *r*(*t*) + *I_const_* + *I_ext_*, where *W* is the connectivity matrix (of size *N* × *N*), 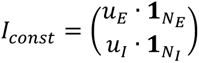 is a constant driving input identical for each neuron class, and *I_ext_* ∈ ℝ*^N^* provides the value of any neuron-specific external (and constant in time) input. The dynamics of neuron *k* evolves according to the differential equation 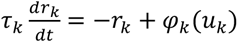 is the transfer (activation) function of the neuron; since we deal with a homogeneous case, it is equal either to *φ_E_* or *φ_I_*, which are the functions for E- and I-neurons. Similarly, *τ_k_* (the neuron’s time constant) is equal either to *τ_E_* or *τ_I_*. The connectivity matrix *W* is generated randomly (Erdos-Renyi case) while respecting Dale principle, i.e., any E-neuron provided only positive inputs, and any I-neuron provided only negative inputs. All E- and I-synapses had equal strength.

The parameters we used for the simulations are as follows.

| Parameter | Value |
| --- | --- |
| Number of E-neurons ($N_E$) | 4000 |
| Number of I-neurons ($N_I$) | 1000 |
| Time constant of E-neurons ($\tau_E$) | 18 ms |
| Time constant of I-neurons ($\tau_I$) | 8 ms |
| Transfer function of E-neurons ($\varphi_E$) | $\varphi_E(x) = [100/(1 + \exp(2 - x)) - 12.5]_+$ |
| Transfer function of I-neurons ($\varphi_I$) | $\varphi_I(x) = [200/(1 + \exp(2 - x)) - 25]_+$ |
| Constant drive of E-neurons ($u_E$) | 25 |
| Constant drive of I-neurons ($u_I$) | 30 |
| Connection probability of $W$ matrix | 0.1 |
| E synaptic strength | 0.0024 |
| I synaptic strength | 0.0088 |
| $I_{ext}$ | 0 for 66.66% of the neurons; otherwise drawn from Gamma(7.5, 1) distribution |

### Conductance-based network model

As with rate network model, we simulated a standard recurrent network of leaky integrate and fire neurons, e.g. (Vogels *et al*., 2005; Brette *et al*., 2007). The spiking network had the same number of excitatory and inhibitory neurons, and the same connection probability as the rate network.

The neurons are standard leaky integrate and fire units. That is, the voltage V(t) of each neuron evolves according to the standard conductance-based leaky integrate-and-fire equation:

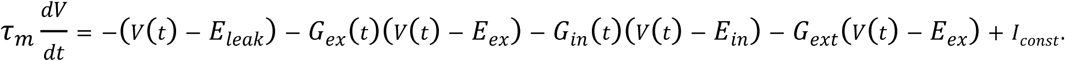

Whenever the voltage reaches the spike threshold (*V_t_*_ℎ_), a spike is emitted, and the membrane potential is instantly clamped at a fixed lower voltage for the duration of the refractory period. Synaptic conductances decay exponentially toward zero in the absence of input and step upward instantaneously upon the arrival of presynaptic spikes.

The parameters we used for the simulations are as follows.

| Parameter | Value |
| --- | --- |
| Number of E-neurons ($N_E$) | 4000 |
| Number of I-neurons ($N_I$) | 1000 |
| Membrane time constant ($\tau_m$) | 20 ms |
| Time constant of E-synapses ($\tau_E$) | 5 ms |
| Time constant of I-synapses ($\tau_I$) | 5 ms |
| E-neuron absolute refractory period ($\tau_{refE}$) | 5 ms |
| I-neuron absolute refractory period ($\tau_{refI}$) | 2 ms |
| Leak reversal potential ($E_{leak}$) | -60 mV |
| Excitatory reversal potential ($E_{ex}$) | 0 mV |
| Inhibitory reversal potential ($E_{in}$) | -80 mV |
| Spiking threshold ($V_{th}$) | -50 mV |
| E-neuron refractory clamping potential | -51.25 mV |
| I-neuron refractory clamping potential | -50.5 mV |
| Connection probability of $W$ matrix | 0.1 |
| E synaptic strength | 0.01 |
| I synaptic strength | 0.06 |
| Constant drive ($I_{const}$) | 12 |
| External input $G_{ext}$ | 0 for 70% of the neurons; otherwise drawn uniformly from the [0, 0.025] interval. |

### Discrimination analysis – single neuron empirical accuracy

For each neuron, we randomly divided the trials into two equal halves (training and test sets, i.e. 2-fold cross-validation). We used the training set to find the optimal threshold *K_t_*_ℎ_ that separates the spike counts of the two conditions. We evaluated the performance of this threshold on the test set of trials.

### Discrimination analysis – single neuron analytical accuracy

Let us first consider the case of Poisson neuron with rates *λ*_1_ and *λ*_2_ in the two conditions (w.l.o.g. *λ*_2_ > *λ*_1_). The probability to observe *k* spikes is given by *P*(*X* = *k*|*λ_i_*) = (*Tλ_i_*)*^k^e*^−*λ*^*^i^*/*k*!, where *T* is trial duration, and *i* = 1,2.

A trial with *k* spikes is classified based on the odds ratio, i.e., *P*(*k*|*cond*_2_)/*P*(*k*|*cond*_1_), a widely-accepted approach both generally and for the concrete case of spike count decoding, e.g., (Jazayeri & Movshon, 2006). Substituting the above expression for Poisson probability mass function shows that this ratio is equal to (*λ*_2_/*λ*_1_)*^k^e*^−*T*(*λ*2−*λ*1)^. It is higher than 1 if and only if *k* >*T*(*λ*_2_−*λ*_1_)/ ln(*λ*_2_/*λ*_1_). This last expression is the threshold value *K_t_*_ℎ_, which provides the optimal way to discriminate between the two conditions given the neuron’s spike count. The predicted discrimination accuracy of the neuron (since the two conditions are equiprobable) is 0.5 · *P*(*X* ≤ *K_t_*_ℎ_|*cond*_1_) + 0.5 · *P*(*X* > *K_t_*_ℎ_|*cond*_2_) = 0.5 · [*F_Pois_* (*K_t_*_ℎ_; *Tλ*_1_) + 1 − *F_Pois_* (*K_t_*_ℎ_; *Tλ*_2_)], where *F_Pois_* is the CDF of the Poisson distribution.

Since most neurons exhibit supra-Poisson variance, the Poisson assumption would lead to an overestimation of the true accuracy. To more accurately model this realistic scenario, we presumed that spike counts *X* have a negative binomial (NB) distribution *NB*(*μ*, *θ*), with mean *μ*, variance *μ* + *μ*^2^/*θ*, and 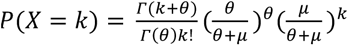. We presume that the Fano factor *F* of each neuron is the same across the two conditions (which is a fairly accurate approximation in our data). It follows that *θ_i_* = *μ_i_*/(*F* − 1), where *μ_i_* is the mean spike count in condition *i* (*i* = 1,2; *μ*_2_ > *μ*_1_).

As before, to classify an observation with *k* spikes, one calculates the odds ratio *P*(*k*|*cond*_2_)/*P*(*k*|*cond*_1_). Using the above expression for NB distribution mass function, and the fact that *μ* /(*θ* + *μ* = (*F* − 1)/*F*, we get: 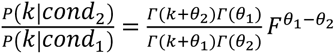.

Unlike the Poisson case, equating this expression to 1 does not lead to a simple, closed form analytical solution. However, it can be solved numerically for each neuron’s set of parameters (*μ*_1_, *μ*_2_ and *F*) to get the threshold value *K_t_*_ℎ_. Once *K_t_*_ℎ_ is found, the predicted discrimination accuracy of the neuron is computed as 0.5 · [*F_NB_*(*K_t_*_ℎ_; *μ*_1_, *θ*_1_) + 1 − *F_NB_*(*K_t_*_ℎ_; *μ*_2_, *θ*_2_)], where *F_NB_* is the CDF of the NB distribution.

### Discrimination analysis – assembly empirical accuracy

For a given assembly (set) of neurons, their spike counts in response to two gratings (across a set of trials) were used as an input to a linear support vector machine (SVM) classification model (MATLAB’s fitcsvm function, with regularisation parameter of 1/30). We used 60 trials for each stimulus, and evaluated the performance using 2-fold cross-validation.

To populate an assembly of a given size (4, 8, 16 or 32, Fig. 6C), neurons were added in the order of their log-rates’ distance from the target log-rate. All neurons from the brain area of interest that were recorded in a session were eligible, provided their log-rate was not closer to another target log-rate. If the number of such neurons was insufficient, the recording session was excluded. If majority of recording sessions did not have enough neurons, the value was not reported (which was the case for sets of size 32 and target mean log-rate of 1.6 = 40 spikes/s).

To populate an assembly with a given budget (e.g. 40 spikes/s, Fig. 6E), neurons were added in the order of their rates’ distance from a target rate. The target rates were the budget value divided by 2*^k^*, *k* = 0, 1, …,6 (to produce assemblies of ∼2*^k^* neurons). All neurons that were recorded in a recording session were eligible, provided their rate was not closer to another target rate. If the number of such neurons was insufficient to reach the target budget, the recording session was excluded. If majority of recording sessions did not have enough neurons, the value was not reported (which was the case for 20 spikes/s budget and target rate of 20/2^6^ = 0.31 spikes/s).

To computationally predict the accuracy of an assembly of N neurons, where each neuron’s activity was specified by its mean log-rate, modulation and Fano factor, we simulated 60 trials for each condition, by randomly drawing the responses of each neuron from NB distribution specified by the three parameters (mean log-rate and modulation are converted into *μ*_1_ and *μ*_2_). We then processed this simulated data in the same day we did for the actual empirical spike counts (described above) to get the predicted accuracy value.

### Analysis code

MATLAB code will be made publicly available upon publication of the manuscript.

## Acknowledgements

This study was supported by the Academy of Medical Sciences and the Wellcome Trust (Springboard award SBF002\1045), BBSRC (grant BB/P020607/1), and Gilgamesh Pharmaceuticals, Inc.

The author used generative AI for some parts of the MATLAB coding and for manuscript editing. All AI output was thoroughly reviewed; the author takes full responsibility for the content of the work.

## Supplementary figures

**Figure S1.**
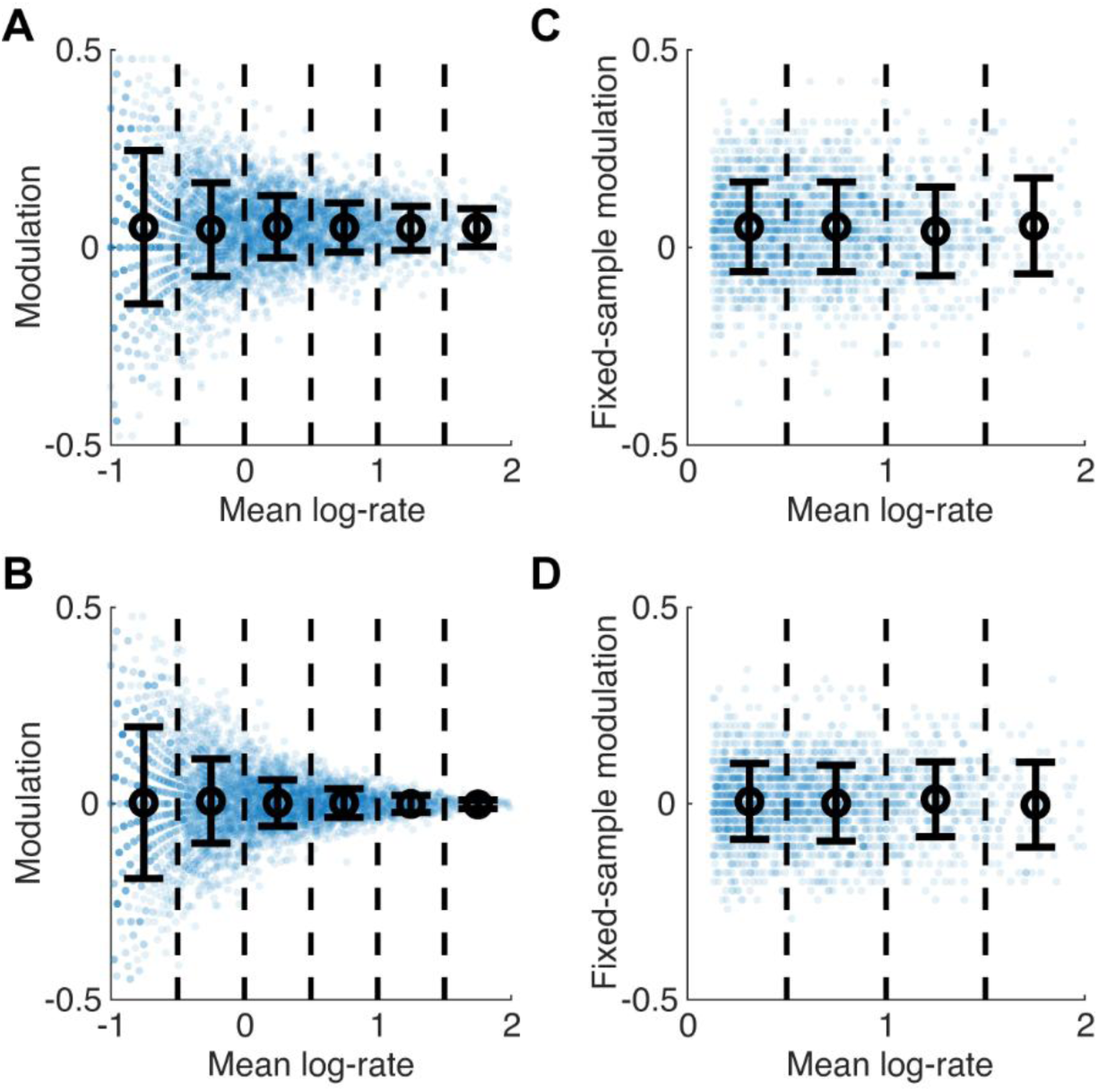
Statistical bias in heteroskedasticity estimation and its removal. **(A)** 5000 Poisson neurons with baseline log-rates between -2 and +2 (corresponding to rates between 0.01 and 100 spikes/s) were simulated. For each neuron, spike counts in 60 trials of 2s in duration were sampled. The baseline log-rate of each neuron was then modulated by adding a value drawn from normal distribution with mean and st.dev. of 0.05, and spike counts for 60 trials with the new log-rate were generated. This data was analysed similarly to the experimental data (cf. Fig. 2F). Heteroskedasticity is clearly seen and is highly statistically significant (p < 10^-100^), however it is the result of estimation bias, since the true modulation mean and variance are independent of the rate. **(B)** Simulation similar to the one in (A) but with no modulation at all – each neuron retains its Poisson rate, so that the two conditions are merely two independent samples of spike counts from Poisson neurons with fixed rate. Again, heteroskedasticity which is caused by sampling bias is clearly seen (p < 10^-100^). **(C)** The data in (A) analysed using fixed-sample modulation approach: whereas the accuracy of the estimate decreased (in particular for fast-firing neurons), it is no longer the case that the estimate is more accurate for fast-firing neurons, and heteroskedasticity is not present (p = 0.79). **(D)** The data in (B) analysed using fixed-sample modulation estimate. Again, heteroskedasticity is not present (p = 0.34).

**Figure S2.**
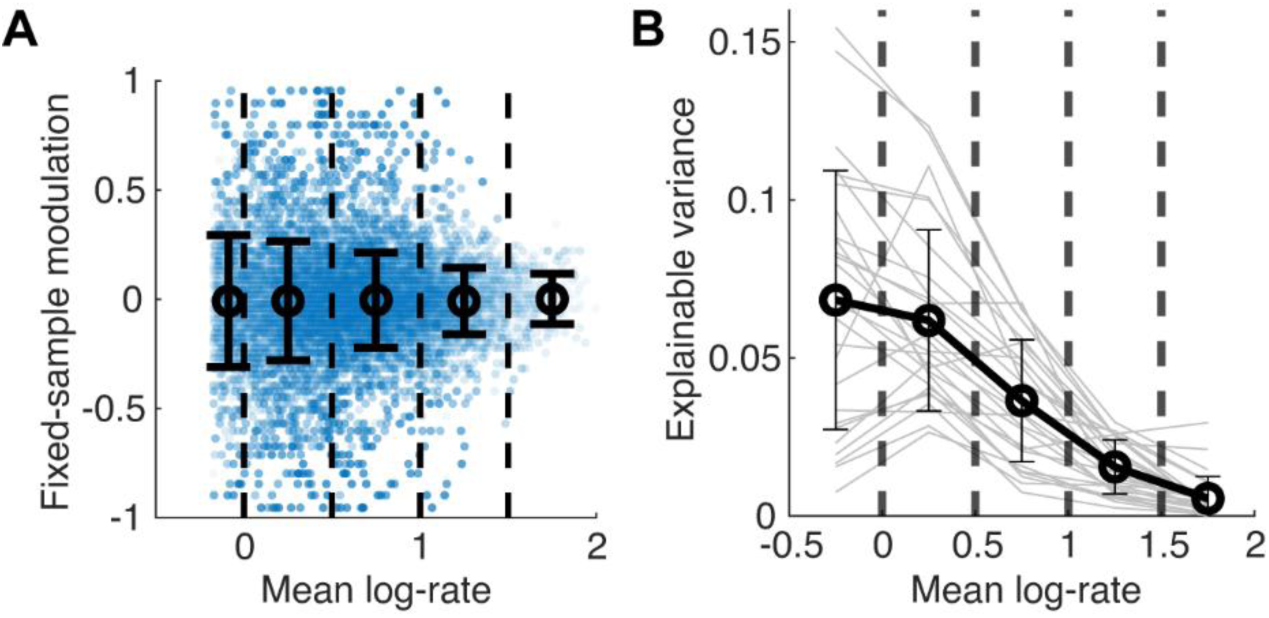
Heteroskedasticity in mouse primary visual cortex. **(A)** Fixed-sample modulation vs mean log-rate (n = 15321 neurons), for 45° vs 90° drifting-grating conditions, showing clear heteroskedasticity (p < 10^-75^). **(B)** Explainable variance in individual sessions and the mean across the sessions. The latter resembles the result with pooled neurons from all sessions (cf. Fig. 3C). Error bars show st. dev. across sessions.

**Figure S3.**
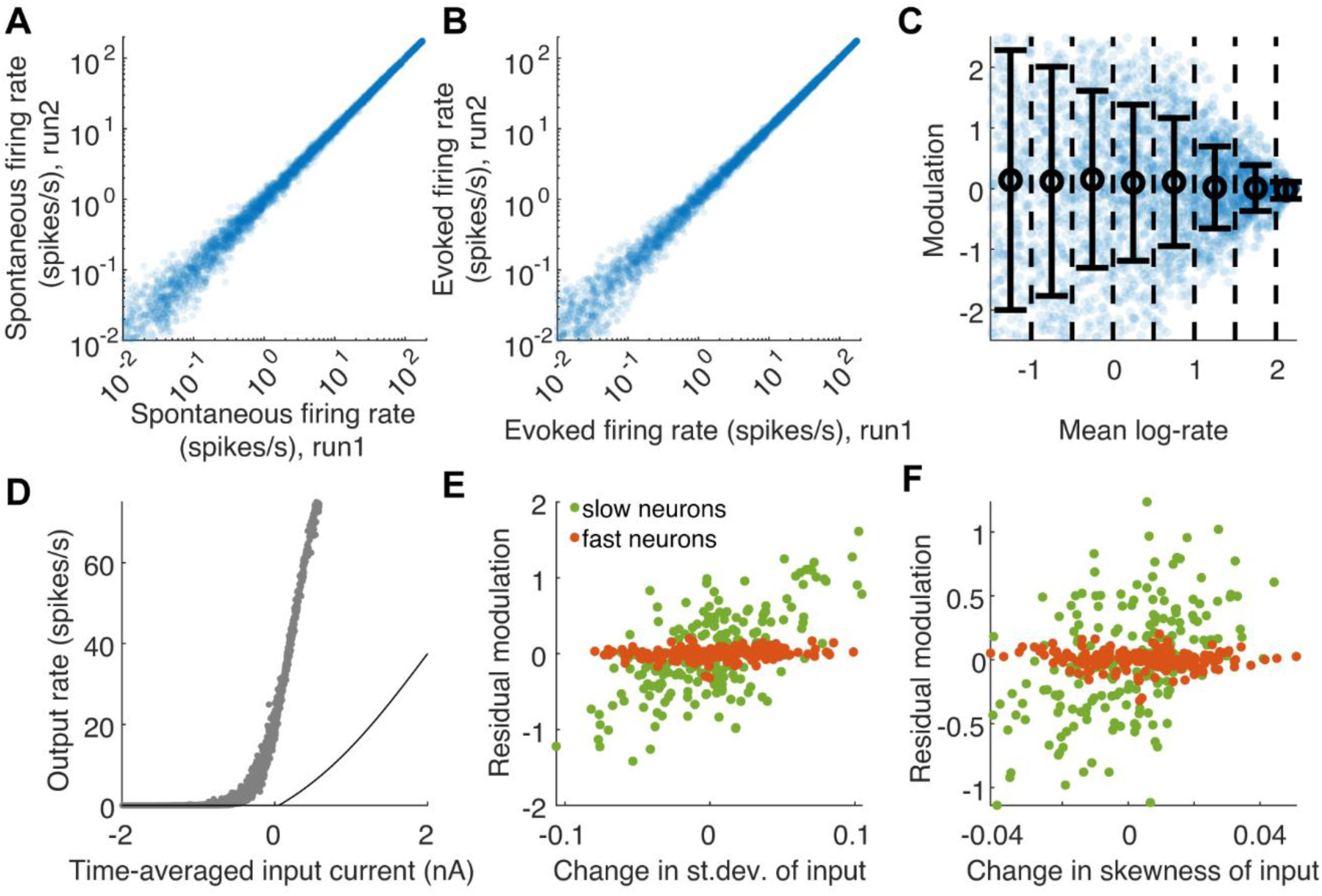
Heteroskedasticity in a rate network model. **(A, B)** Rates of neurons in two independent runs of the model network, plotted against each other (in spontaneous and evoked conditions). The rates are highly consistent, indicating that the difference in rate that we observed across conditions (cf. Fig. 5B) was caused by change in the state of the network. Some heteroskedasticity is visibly present, indicating true higher within-condition rate variability of slow neurons. (Note that in the case of a rate model the activity of each neuron is a continuous signal rather than a spike train; in particular, the fixed-sample modulation estimate, which works with discrete spikes, is not applicable). **(C)** Heteroskedasticity in the model was present not only for changes between spontaneous and evoked conditions, but also between a pair of evoked conditions (p < 10^-100^) when the parameters of the external stimulus were sampled independently. **(D)** The data from Fig. 5E replotted together with the activation function of E-neurons in the model (black curve). The major discrepancy between the two highlights the major role of the network dynamics (fluctuations in the input current) in shaping the effective F-I relationship in the network. **(E)** Change in the st.dev. of the time-averaged input current caused by shift between spontaneous and evoked regimes has different modulation gain in slow- and fast-firing E-neurons. Y-axis shows the residual modulation, after subtracting the prediction based on linear regression on the mean time-averaged input (Fig. 5D). **(F)** Change in the skewness of the time-averaged input current caused by shift between spontaneous and evoked regimes has different modulation gain in slow and fast firing E-neurons. Y-axis shows the residual modulation, after subtracting the prediction based on linear regression on the mean and st.dev. of the time-averaged input.

**Figure S4.**
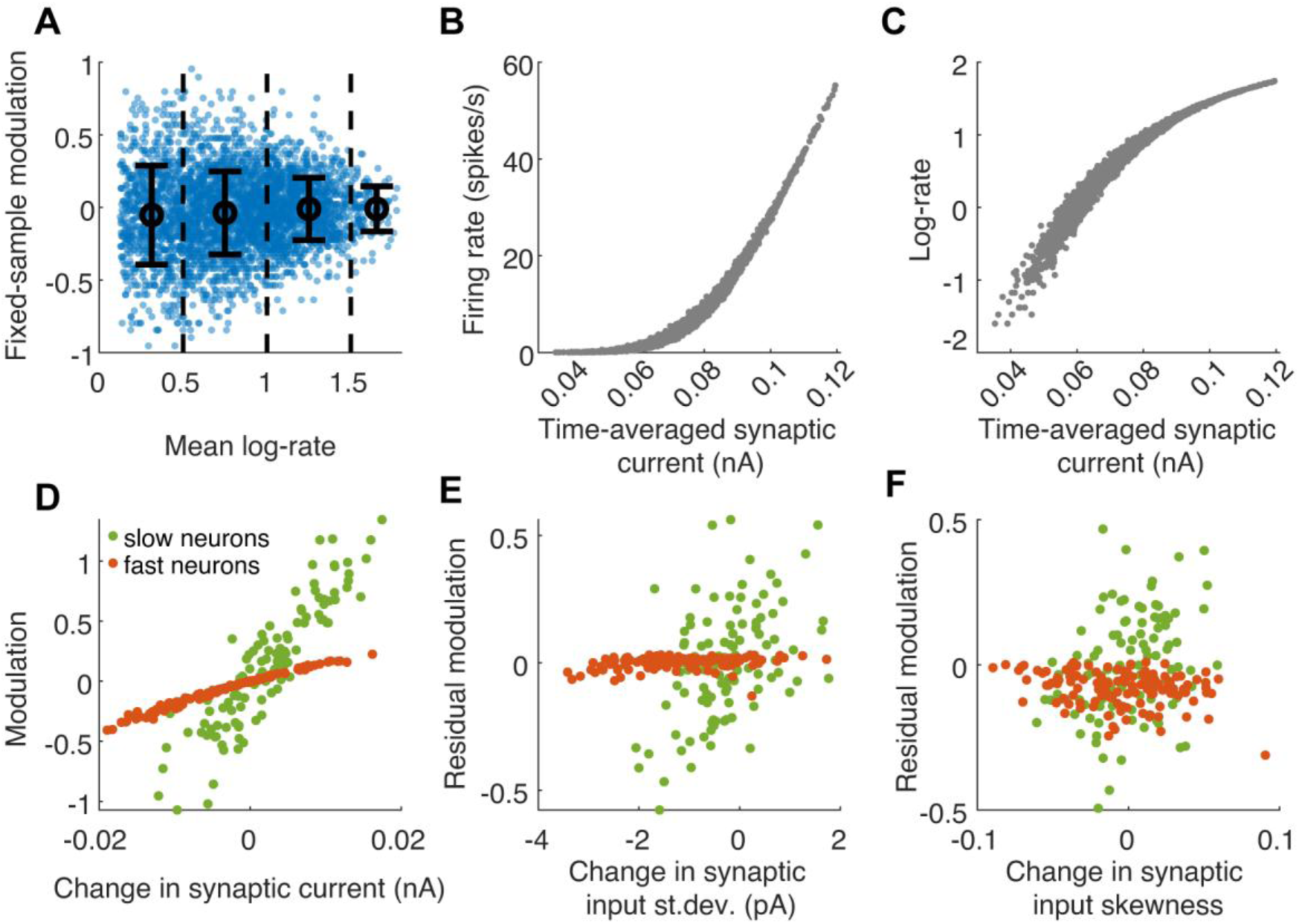
Heteroskedasticity in a conductance based, spiking network model. **(A)** Heteroskedasticity for a change between two evoked regimes (p < 10^-30^; cf. Fig. 5C). **(B)** The relationship between time-averaged input current and output rate for E-neurons, notice its convex shape (cf. Fig. 5E). **(C)** As in (B), with output rates shown on log-scale, note the concave shape (cf. Fig. 5F). **(D)** Change in the time-averaged input current caused by shift between the two regimes has different modulation gain in slow-firing (time-averaged input is 0.04-0.05 nA) and fast-firing (time-averaged input is 0.10-0.11 nA) E-neurons (cf. Fig. 5D). **(E)** Change in the st.dev. of the input caused by shift between the two regimes has different modulation gain in slow- and fast-firing E-neurons (cf. Fig. S3E). Y-axis shows the residual modulation, after subtracting the prediction based on linear regression on the time-averaged input (D). For slow neurons the correlation between change in st.dev. and the residual modulation is significant (r = 0.34, p = 3.7×10^-4^; Pearson correlation), for fast-firing neurons this is not the case (p = 0.25). **(F)** Change in the skewness of the input caused by shift between the two regimes has different modulation gain in slow- and fast-firing E-neurons (cf. Fig. S3F). Y-axis shows the residual modulation, after subtracting the prediction based on linear regression on the mean and st.dev. of the input (D, E). For slow neurons the correlation between change in skewness and the residual modulation is significant (r = 0.24, p = 0.013; Pearson correlation), for fast-firing neurons this is not the case (p = 0.72).

**Figure S5.**
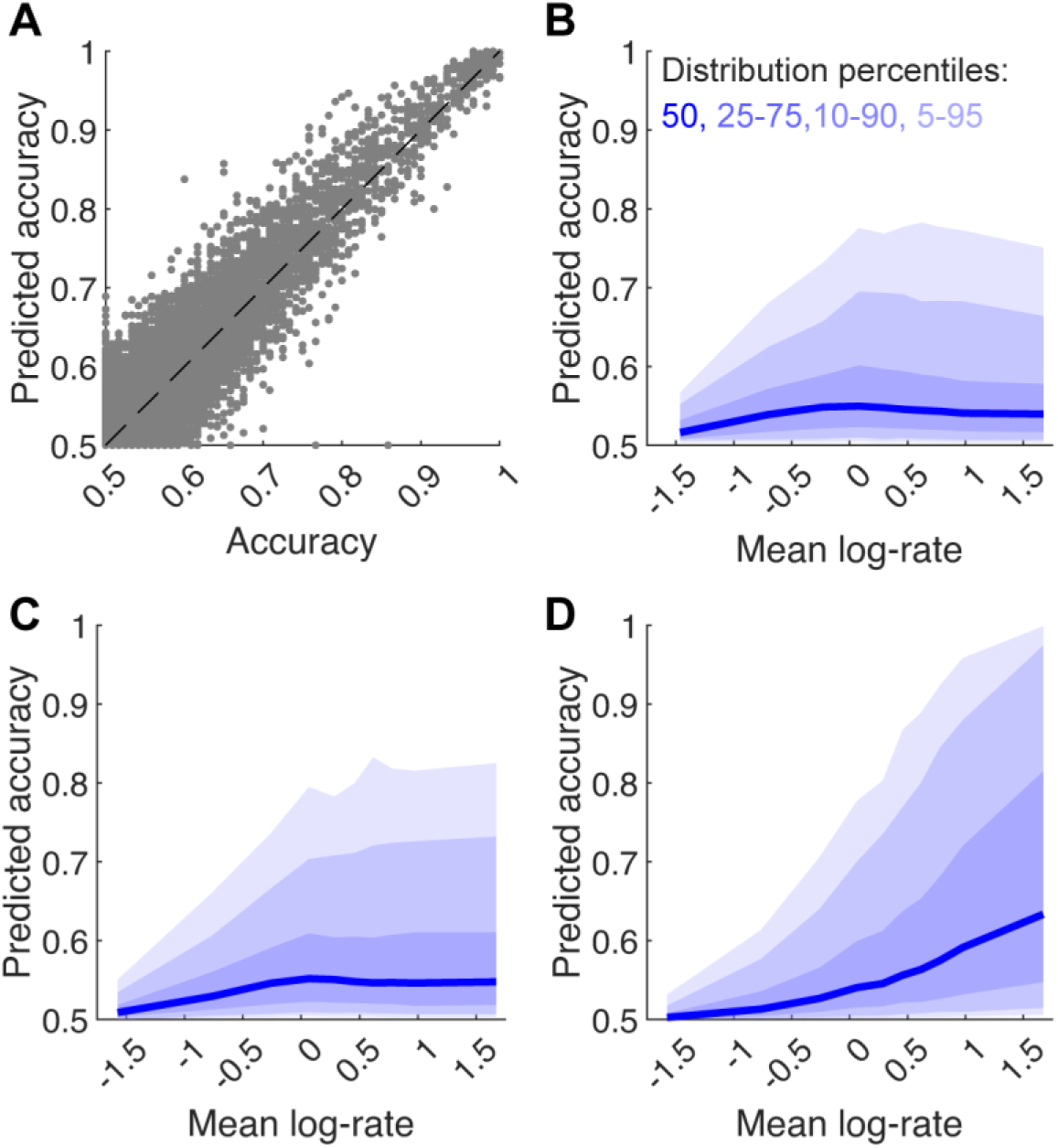
Single neuron discrimination accuracy. **(A)** Discrimination accuracy based on spike counts of single visual cortex neurons can be either directly estimated from the 60 trials data in each condition (x-axis), or analytically predicted from just three parameters: mean log-rate, modulation and Fano factor of the neuron (y-axis). The analytical prediction closely matches the direct estimate (R^2^ = 0.81, p < 10^-100^). **(B)** The distribution of analytically predicted accuracy vs mean log-rate closely matches the distribution of empirical accuracy (cf. Fig. 6A). **(C, D)** The analytical prediction allows to examine how the accuracy distribution would have looked like if Fano factor (C) or both Fano factor and modulation (D) were independent of mean log-rate (achieved by shuffling these parameters across the neuronal population). This shows that the empirical distribution (shown in Fig. 6A, and in B) is primarily shaped by the dependence of modulation on rate, whereas Fano factor’s dependence on rate has a marginal effect (see also Fig. 6B).

**Figure S6.**
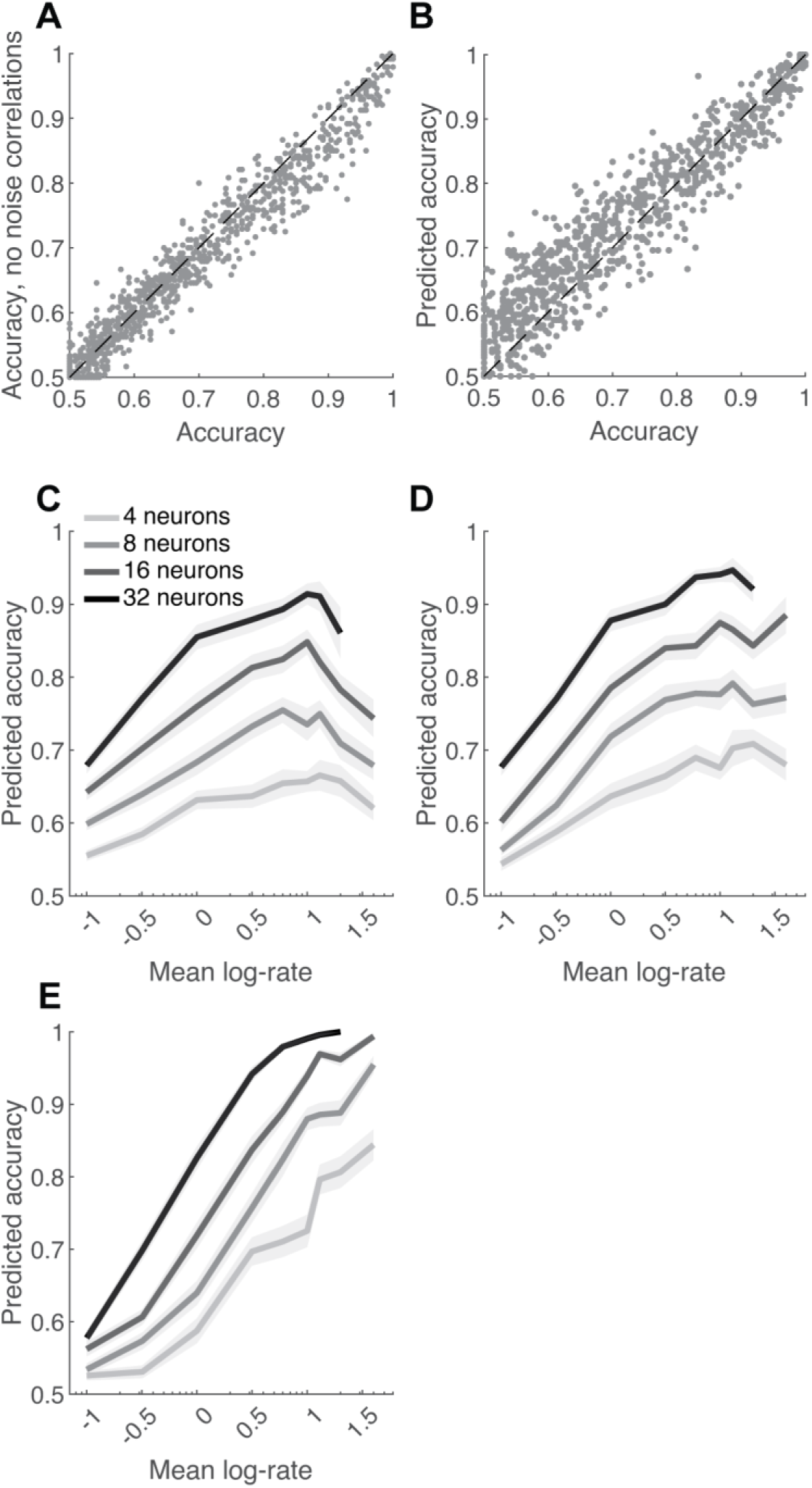
Discrimination accuracy of a reader with input wiring constraint. We formed assemblies of size 4, 8, 16, and 32 from simultaneously recorded visual cortex neurons with similar rate. **(A)** The discrimination accuracy of these assemblies was only weakly dependent on noise correlations, since trial shuffling did not significantly alter accuracy (R^2^ = 0.94, p < 10^-100^, n = 865 assemblies from 26 recordings). **(B)** An assembly’s predicted accuracy, which was calculated based on the mean log-rate, modulation and Fano factor of each neuron (and presuming the neurons are independent) closely matched the empirical accuracy values (R^2^ = 0.86, P < 10^-100^). **(C)** Discrimination accuracy using spike counts from randomly selected assemblies of neurons with matching mean log-rates spanning the rate spectrum. The accuracy was predicted, but the results are similar to Fig. 6C. **(D)** Discrimination accuracy of assemblies in a synthetic scenario in which Fano factor is independent of the rate of the neurons (created by shuffling the Fano factor parameter across the neuronal population). Note the similarity to (C). **(E)** Discrimination accuracy of assemblies in a synthetic scenario in which modulation and Fano factor are independent of the rate of the neurons (created by shuffling both parameters across the neuronal population). Note the difference vs (C, D) and similarity to Fig. 6D.

**Figure S7.**
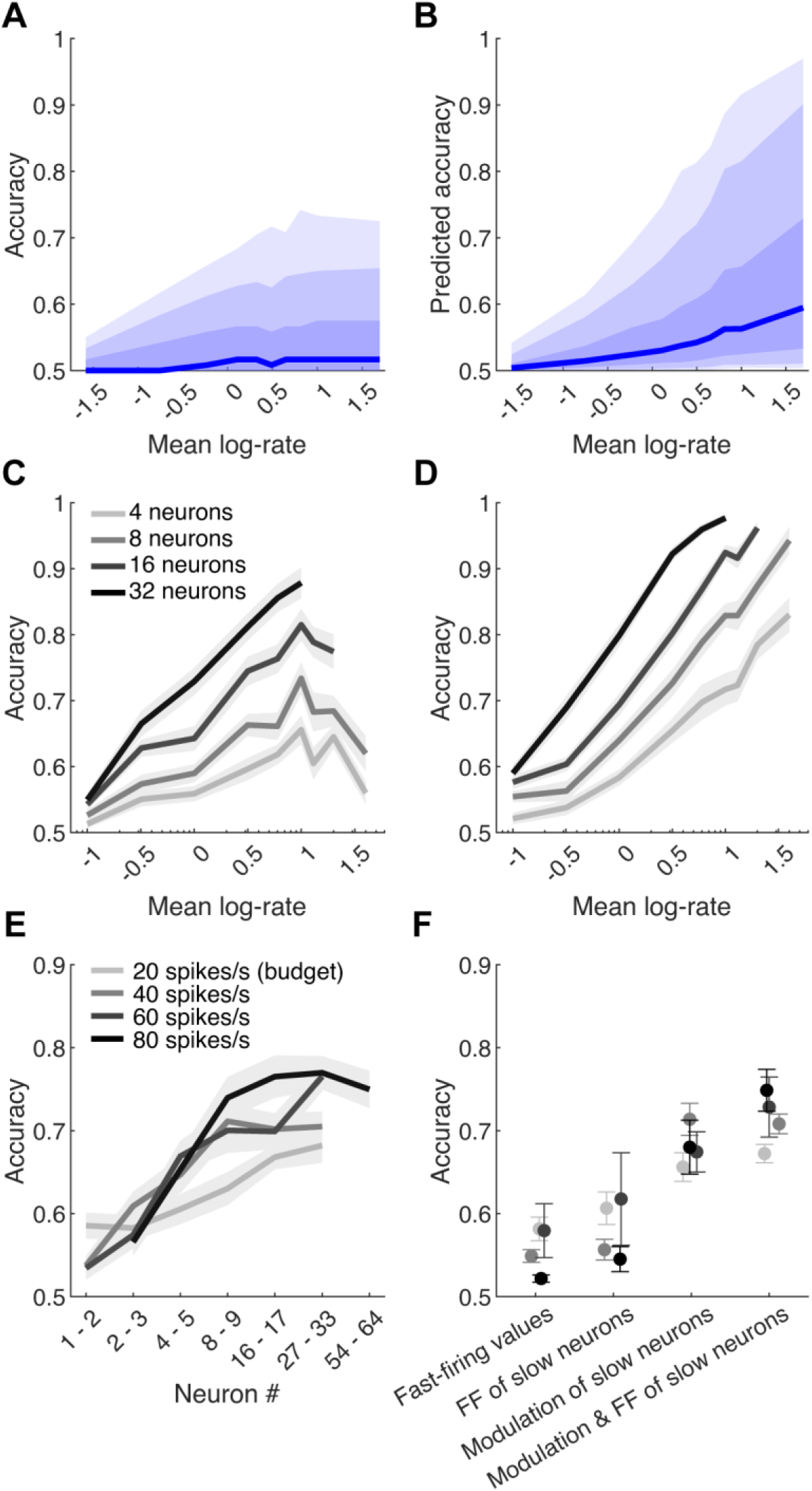
Heteroskedasticity’s impact on readout accuracy in CA1. Same format as Fig. 6, for neurons and assemblies in the CA1.

**Figure S8.**
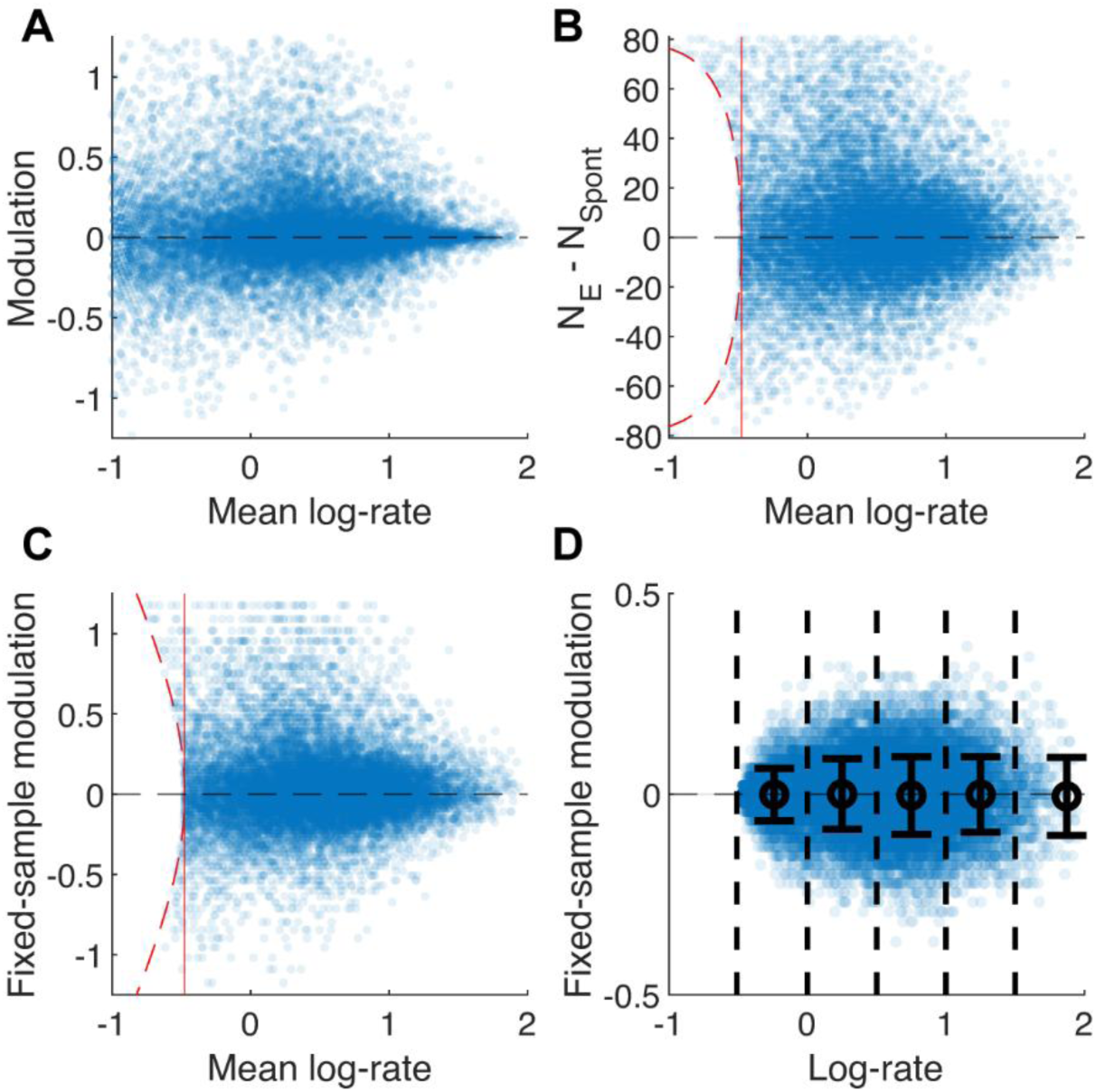
Fixed-sample modulation estimate. **(A)** Modulation vs mean log-rate for spontaneous and evoked conditions (as in Fig. 2E). **(B)** Subsampling leaves exactly *N_T_* = 80 randomly selected spikes for each neuron, and neurons with less than *N_T_* spikes (points to the left of the dashed red curve) are removed from the analysis. *N_E_* and *N_Spont_* denote the total number of spikes in each condition after subsampling (*N_E_* + *N_Spont_* = *N_T_*). **(C)** As in (B), with y-axis showing the fixed-sample modulation, defined as log_10_(*N_E_* /*N_Spont_*). Neurons falling between the red dashed curve and the red vertical line marking its apex are also removed, in order not to bias the variance analysis which is based on binning of the points by their mean log-rate. **(D)** Demonstration of heteroskedasticity introduced by the subsampling procedure itself (in the case of identical spike counts in the two conditions). Importantly, it is in reverse direction from the one seen in the data.

## References

Ahmadian Y, Rubin DB & Miller KD (2013). Analysis of the Stabilized Supralinear Network. Neural Comput 25, 1994–2037.

Anderson JS, Lampl I, Gillespie DC & Ferster D (2000). The Contribution of Noise to Contrast Invariance of Orientation Tuning in Cat Visual Cortex. Science 290, 1968– 1972.

Brecier A, Borel M, Urbain N & Gentet LJ (2022). Vigilance and Behavioral State-Dependent Modulation of Cortical Neuronal Activity throughout the Sleep/Wake Cycle. J Neurosci 42, 4852–4866.

Brette R et al. (2007). Simulation of networks of spiking neurons: a review of tools and strategies. J Comput Neurosci 23, 349–398.

Brunel N (2000). Dynamics of networks of randomly connected excitatory and inhibitory spiking neurons. J Physiol Paris 94, 445–463.

Bruno RM & Sakmann B (2006). Cortex is driven by weak but synchronously active thalamocortical synapses. Science 312, 1622–1627.

Buzsaki G & Mizuseki K (2014). The log-dynamic brain: how skewed distributions affect network operations. Nat Rev Neurosci 15, 264–278.

Chance FS, Abbott LF & Reyes AD (2002). Gain modulation from background synaptic input. Neuron 35, 773–782.

Chen Y, Kloos M, Varga Z, Zhang Y, Piro I, Sato TK, Sakmann B, Nelken I & Konnerth A (2026). Thalamic activation of the visual cortex at the single-synapse level. Science 391, 1349–1354.

Dayan P & Abbott LF (2005). Theoretical Neuroscience: Computational and Mathematical Modeling of Neural Systems, Revised ed. edition. The MIT Press, Cambridge, Mass.

Dearnley B, Dervinis M, Shaw M & Okun M (2021). Stretching and squeezing of neuronal log firing rate distribution by psychedelic and intrinsic brain state transitions. bioRxiv 2021.08.22.457198.

Dearnley B, Jones M, Dervinis M & Okun M (2023). Brain state transitions primarily impact the spontaneous rate of slow-firing neurons. Cell Reports 42, 113185.

Gentet LJ, Kremer Y, Taniguchi H, Huang ZJ, Staiger JF & Petersen CCH (2012). Unique functional properties of somatostatin-expressing GABAergic neurons in mouse barrel cortex. Nat Neurosci; DOI: 10.1038/nn.3051.

Gerstner W, Kistler WM, Naud R & Paninski L (2014). Neuronal Dynamics: From Single Neurons to Networks and Models of Cognition, 1st edition. Cambridge University Press.

Goris RLT, Movshon JA & Simoncelli EP (2014). Partitioning neuronal variability. Nat Neurosci 17, 858–865.

Graf ABA, Kohn A, Jazayeri M & Movshon JA (2011). Decoding the activity of neuronal populations in macaque primary visual cortex. Nat Neurosci 14, 239–245.

Hansel D & Vreeswijk C van (2002). How Noise Contributes to Contrast Invariance of Orientation Tuning in Cat Visual Cortex. J Neurosci 22, 5118–5128.

James G, Witten D, Hastie T & Tibshirani R (2013). An Introduction to Statistical Learning: with Applications in R: 103, 1st ed. 2013, Corr. 7th printing 2017 edition. Springer, New York.

Jazayeri M & Movshon JA (2006). Optimal representation of sensory information by neural populations. Nature neuroscience 9, 690–696.

Jones M (2024). Structure of modulation of neuronal firing rates across diflerent brain state transitions. (PhD Thesis thesis). University of Leicester. Available at: https://figshare.le.ac.uk/articles/thesis/Structure_of_modulation_of_neuronal_firing_rates_across_different_brain_state_transitions_/27889815 [Accessed March 4, 2025].

Jun JJ et al. (2017). Fully integrated silicon probes for high-density recording of neural activity. Nature 551, 232–236.

Lawlor PN, Perich MG, Miller LE & Kording KP (2018). Linear-nonlinear-time-warp-poisson models of neural activity. J Comput Neurosci 45, 173–191.

Levenstein D & Okun M (2023). Logarithmically scaled, gamma distributed neuronal spiking. The Journal of Physiology 601, 3055–3069.

Levenstein D, Watson BO, Rinzel J & Buzsáki G (2017). Sleep regulation of the distribution of cortical firing rates. Current Opinion in Neurobiology 44, 34–42.

McCormick DA, Connors BW, Lighthall JW & Prince DA (1985). Comparative electrophysiology of pyramidal and sparsely spiny stellate neurons of the neocortex. J Neurophysiol 54, 782–806.

Miller KD & Troyer TW (2002). Neural Noise Can Explain Expansive, Power-Law Nonlinearities in Neural Response Functions. Journal of Neurophysiology 87, 653– 659.

O’Connor DH, Peron SP, Huber D & Svoboda K (2010). Neural activity in barrel cortex underlying vibrissa-based object localization in mice. Neuron 67, 1048–1061.

Perich MG, Lawlor PN, Kording KP & Miller LE (2018). Extracellular neural recordings from macaque primary and dorsal premotor motor cortex during a sequential reaching task. CRCNS.org.; DOI: 10.6080/K0FT8J72. Available at: crcns.org/data-sets/motor-cortex/pmd-1/about-pmd-1.

Petersen PC & Berg RW (2016). Lognormal firing rate distribution reveals prominent fluctuation–driven regime in spinal motor networks. eLife 5, e18805.

Priebe NJ & Ferster D (2008). Inhibition, spike threshold, and stimulus selectivity in primary visual cortex. Neuron 57, 482–497.

Resulaj A, Ruediger S, Olsen SR & Scanziani M (2018). First spikes in visual cortex enable perceptual discrimination. Elife 7, e34044.

Roxin A, Brunel N, Hansel D, Mongillo G & Vreeswijk C van (2011). On the Distribution of Firing Rates in Networks of Cortical Neurons. J Neurosci 31, 16217–16226.

Sanzeni A, Palmigiano A, Nguyen TH, Luo J, Nassi JJ, Reynolds JH, Histed MH, Miller KD & Brunel N (2023). Mechanisms underlying reshuffling of visual responses by optogenetic stimulation in mice and monkeys. Neuron 111, 4102–4115.e9.

Siegle JH et al. (2021). Survey of spiking in the mouse visual system reveals functional hierarchy. Nature 592, 86–92.

Silver RA (2010). Neuronal arithmetic. Nat Rev Neurosci 11, 474–489.

Sterling P & Laughlin S (2015). Principles of Neural Design. The MIT Press.

Stringer C, Michaelos M, Tsyboulski D, Lindo SE & Pachitariu M (2021). High-precision coding in visual cortex. Cell 184, 2767–2778.e15.

Tasic B et al. (2018). Shared and distinct transcriptomic cell types across neocortical areas. Nature 563, 72–78.

Trepka EB, Zhu S, Xia R, Chen X & Moore T (2022). Functional interactions among neurons within single columns of macaque V1 ed. Vinck M, Behrens TE & Kremkow J. eLife 11, e79322.

Trojanowski NF, Bottorff J & Turrigiano GG (2021). Activity labeling in vivo using CaMPARI2 reveals intrinsic and synaptic differences between neurons with high and low firing rate set points. Neuron 109, 663–676.e5.

Valero M, Abad-Perez P, Gallardo A, Picco M, García-Hernandez R, Brotons J, Martínez-Félix A, Machold R, Rudy B & Buzsaki G (2025). Cooperative actions of interneuron families support the hippocampal spatial code. Science 389, eadv5638.

Vogels TP, Rajan K & Abbott LF (2005). Neural network dynamics. Annu Rev Neurosci 28, 357–376.

Yao S et al. (2023). A whole-brain monosynaptic input connectome to neuron classes in mouse visual cortex. Nat Neurosci 26, 350–364.

Yassin L, Benedetti BL, Jouhanneau J-S, Wen JA, Poulet JFA & Barth AL (2010). An embedded subnetwork of highly active neurons in the neocortex. Neuron 68, 1043– 1050.

Zhang M, Pan X, Jung W, Halpern AR, Eichhorn SW, Lei Z, Cohen L, Smith KA, Tasic B, Yao Z, Zeng H & Zhuang X (2023). Molecularly defined and spatially resolved cell atlas of the whole mouse brain. Nature 624, 343–354.

Zhu S, Xia R, Chen X & Moore T (2020). Heterogeneity of Neuronal Populations Within Columns of Primate V1 Revealed by High-Density Recordings. bioRxiv 2020.12.22.424048.

